# Universal trends of post-duplication evolution revealed by the genomes of 13 *Paramecium* species sharing an ancestral whole-genome duplication

**DOI:** 10.1101/573576

**Authors:** Jean-Francois Gout, Parul Johri, Olivier Arnaiz, Thomas G. Doak, Simran Bhullar, Arnaud Couloux, Fréderic Guérin, Sophie Malinsky, Linda Sperling, Karine Labadie, Eric Meyer, Sandra Duharcourt, Michael Lynch

**Affiliations:** Department of Biology, Indiana University, Bloomington, IN, USA; School of Life Sciences, Biodesign Center for Mechanisms of Evolution, Arizona State University, Tempe, Arizona, USA; Department of Biological Sciences, Mississippi State University, PO BOX GY, Mississippi State, MS 39762, USA; Institute for Integrative Biology of the Cell (I2BC), CEA, CNRS, Univ. Paris-Sud, Universite Paris-Saclay, 91198 Gif-sur-Yvette cedex, France; National Center for Genome Analysis Support, Indiana University, Bloomington, IN 47405; Genoscope/Centre National de Sequençage, 2 rue Gaston Cremieux, CP 5706, 91057 Evry CEDEX, France; Institut Jacques Monod, CNRS, UMR 7592, Université Paris Diderot, Sorbonne Paris Cité, Paris, F-75205 France; IBENS, Département de Biologie, Ecole Normale Supérieure, CNRS, Inserm, PSL Research University, F-75005 Paris, France

## Abstract

Whole-Genome Duplications (WGDs) have shaped the gene repertoire of many eukaryotic lineages. The redundancy created by WGDs typically results in a phase of massive gene loss. However, some WGD-derived paralogs are maintained over long evolutionary periods and the relative contributions of different selective pressures to their maintenance is still debated. Previous studies have revealed a history of three successive WGDs in the lineage of the ciliate *Paramecium tetraurelia* and two of its sister species from the *P. aurelia* complex. Here, we report the genome sequence and analysis of 10 additional *P. aurelia* species and one additional outgroup, allowing us to track post-WGD evolution in 13 species that share a common ancestral WGD. We found similar biases in gene retention compatible with dosage constraints playing a major role opposing post-WGD gene loss across all 13 species. Interestingly we found that post-WGD gene loss was slower in *Paramecium* than in other species having experienced genome duplication, suggesting that the selective pressures against post-WGD gene loss are especially strong in *Paramecium*. We also report a lack of recent segmental duplications in *Paramecium*, which we interpret as additional evidence for strong selective pressures against individual genes dosage changes. Finally, we hope that this exceptional dataset of 13 species sharing an ancestral WGD and two closely related outgroup species will be a useful resource for future studies and will help establish *Paramecium* as a major model organism in the study of post-WGD evolution.

## Introduction

Gene duplication is a common type of mutation that can occur at frequencies rivaling that of point mutations (Lynch 2007; Lipinski, et al. 2011; Schrider, et al. 2013; Reams and Roth 2015). Because duplicated genes are often redundant, mutations crippling one copy are expected to frequently drift to fixation, unaffected by selection. As a consequence, the fate of most duplicated genes is to rapidly turn into pseudogenes and eventually evolve beyond recognition. However, ancient duplicated genes are ubiquitous in the genomes of all free-living organisms sequenced to date (Zhang 2003), to the point that almost all genes in the human genome might be the result of an ancient gene duplication (Britten 2006). Therefore, selective pressures opposing the loss of duplicated genes must be commonly operating despite the initial redundancy between the two copies.

Several models have been proposed to explain the long term retention of duplicated genes. Retention can happen through change in function when one copy acquires beneficial mutations conferring a new function (neofunctionalization (Ohno 1970)) or when each copy independently loses a subset of the functions performed by the ancestral (pre-duplication) gene (subfunctionalization (Force, et al. 1999; Lynch and Force 2000)). It should also be noted that new mutations may not always be necessary to promote gene retention through change of function. Indeed, segmental duplications can lead to partial gene duplications so that the new copy already diverges from the ancestral sequence immediately following the duplication event. Even when all of the exons of a gene are duplicated, regulatory regions and chromatin context might be different between the two copies, potentially resulting in the new copy being born with a different expression profile. Finally, duplicated genes can also be retained without a change in their function. Dosage constraints can drive selection to maintain duplicated genes performing the same function when multiple copies are needed to produce the required amount of transcripts (Edger and Pires 2009; Birchler and Veitia 2012). In this case, selection is expected to act on the total (*i.e.* combined between all copies) amount of transcripts produced.

In its most extreme form, duplication can encompass the entire genome, creating a new copy of each gene. Such Whole-Genome Duplication (WGD) events are not uncommon, with evidence of ancient WGDs in the lineages of many eukaryotes including the budding yeast (Kellis, et al. 2004), insects (Li, et al. 2018), the African clawed frog (Session, et al. 2016), the rainbow trout (Berthelot, et al. 2014) and *Paramecium* (Aury, et al. 2006). It is also now widely accepted that two successive rounds of WGDs occurred in the ancestor of vertebrates (Hokamp, et al. 2003; Dehal and Boore 2005; Holland and Ocampo Daza 2018) and an additional round of genome duplication in the lineage leading to all teleost fish (Meyer and Schartl 1999; Jaillon, et al. 2004; Howe, et al. 2013; Glasauer and Neuhauss 2014). Additionally, WGDs are extremely common in land plants, to the point that all angiosperms are believed to have experienced at least one round of genome duplication in their history (De Bodt, et al. 2005; Ren, et al. 2018). Because they create the opportunity for thousands of genes to evolve new functions, WGDs have been suggested to be responsible for the evolutionary success of several lineages (De Bodt, et al. 2005; Glasauer and Neuhauss 2014). However, the precise link between WGDs and evolutionary success remains unclear (Clarke, et al. 2016; Laurent, et al. 2017).

Here, we investigate the evolutionary trajectories of duplicated genes across multiple *Paramecium* species that share a common ancestral WGD. The initial sequencing of the *Paramecium tetraurelia* genome revealed a history of three (possibly four) successive WGDs (Aury, et al. 2006). Similarly to what was observed in other lineages having experienced WGDs, the *Paramecium* WGD was followed by a phase of massive gene loss and only a fraction of WGD-derived paralogs (ohnologs) have been retained in two copies. Still, about 50% of ohnologs from the most recent WGD are retained in two copies in *P. tetraurelia* (Aury, et al. 2006), a situation very different from that in the budding yeast (about 10% retention rate (Scannell, et al. 2007)), the other widely studied unicellular eukaryote with an ancestral WGD. This situation makes *Paramecium* an ideal model organism for studying the earlier stages of post-WGD genome evolution. Interestingly, the most recent *Paramecium* WGD shortly predates the first speciation events in the formation of the *P. aurelia* group of 15 cryptic *Paramecium* species (Aury, et al. 2006; McGrath, Gout, Doak, et al. 2014; McGrath, Gout, Johri, et al. 2014). It has been suggested that reciprocal gene losses following genome duplication have fueled the speciation of *P. aurelias* (Aury, et al. 2006; McGrath, Gout, Johri, et al. 2014). However, it seems unlikely that this WGD resulted in massive genetic and phenotypic innovations as suggested for yeast (Huminiecki and Conant 2012), plants (Van de Peer, et al. 2009; Edger, et al. 2015) and vertebrates (Voldoire, et al. 2017; Clark and Donoghue 2018). Indeed, the 15 species in the *P. aurelia* complex are morphologically so similar to each other that they were thought to be one single species (*Paramecium aurelia*) until Tracy Sonneborn discovered the existence of mating types and realized that the species he was studying, *P. aurelia,* was in fact a complex of many genetically isolated species (Sonneborn 1937). This apparent lack of post-WGD phenotypic innovation is also supported by a recent study focusing on the Rab GTPase gene family which found that the recent ohnologs in this gene family did not show any sign of functional diversification (Bright, et al. 2017). These observations suggest that neofunctionalization probably did not play a major role in the retention of ohnologs following the most recent genome duplication in *Paramecium*. Our previous studies based on three *P. aurelia* genomes and one pre-WGD outgroup also pointed to an important role of dosage constraint in the retention pattern of ohnologs in *Paramecium* (McGrath, Gout, Doak, et al. 2014; McGrath, Gout, Johri, et al. 2014; Gout and Lynch 2015).

Here, we sought to increase our understanding of post-WGD genome evolution by sequencing the somatic (macronucleus) genomes of the remaining *P. aurelia* species and mapping the trajectories of all genes created by the recent WGD and the subsequent speciation events in *P. aurelia*. We also generated transcriptomic data for each species in order to characterize gene expression levels and better understand the role of expression level and dosage constraints in ohnolog retention. With this dataset, we aim at providing the scientific community with resources comparable to those available in the budding yeast (Byrne and Wolfe 2005) and help establishing *Paramecium* as another model organism for the study of post-WGD genome evolution.

## Results and Discussion

### Genome and transcriptome sequence of 13 *P. aurelia* species that share a common ancestral Whole-Genome Duplication

Previous studies have revealed a history of Whole-Genome Duplication (WGDs) in the lineage of *Paramecium* species belonging to the *P. aurelia* complex (Aury, et al. 2006; McGrath, Gout, Doak, et al. 2014; McGrath, Gout, Johri, et al. 2014), a group of species thought to have speciated shortly following the most recent *Paramecium* WGD. To further understand the evolutionary trajectories of WGD-derived paralogs (ohnologs) we sought to complete our previous efforts (McGrath, Gout, Doak, et al. 2014; McGrath, Gout, Johri, et al. 2014) and sequenced the remaining species from the *P. aurelia* group as well as an additional closely related outgroup. The complete dataset contains 13 species from the *P. aurelia* group and the two outgroups that diverged before the most recent WGD: *P. caudatum* and *P. multimicronucleatum*. All genomes (including previously published ones) were annotated using the EuGene pipeline (Foissac 2008; Arnaiz, et al. 2017) and evidence for a recent WGD was observed in all species of the *P. aurelia* group but absent from both outgroup species (methods). The fraction of ohnologous pairs that maintained both genes intact varied from 0.39 (*P. tredecaurelia*) to 0.58 (*P. jeningsi*) with a median retention rate across species of 0.52 (Table S1).

### Phylogeny of the *P. aurelia* complex

In order to investigate the fate of duplicated genes, we mapped all orthologous and paralogous relationships in the *P. aurelia* complex. Because the first speciation events occurred very shortly after the genome duplication, discriminating orthologs from paralogos when analyzing the most divergent *P. aurelia* species is challenging. We used PoFF (Lechner, et al. 2014) to infer orthology relationships and took advantage of the low rate of large-scale genomic rearrangements in *Paramecium* to assign orthology by blocs of conserved syntheny (Methods). Assigning orthology relationships in blocs of genes gives us more phylogenetic signal for each orthology assignment and increases our capacity to accurately discriminate orthologs from paralogs in deep species comparisons. The final orthology assignments were used to build a reliable phylogeny of the *P. aurelia* group (Figure 1). All positions are strongly supported (100%) by bootsraping, with the exception of *P. biaurelia* (60%).

**Figure 1.**
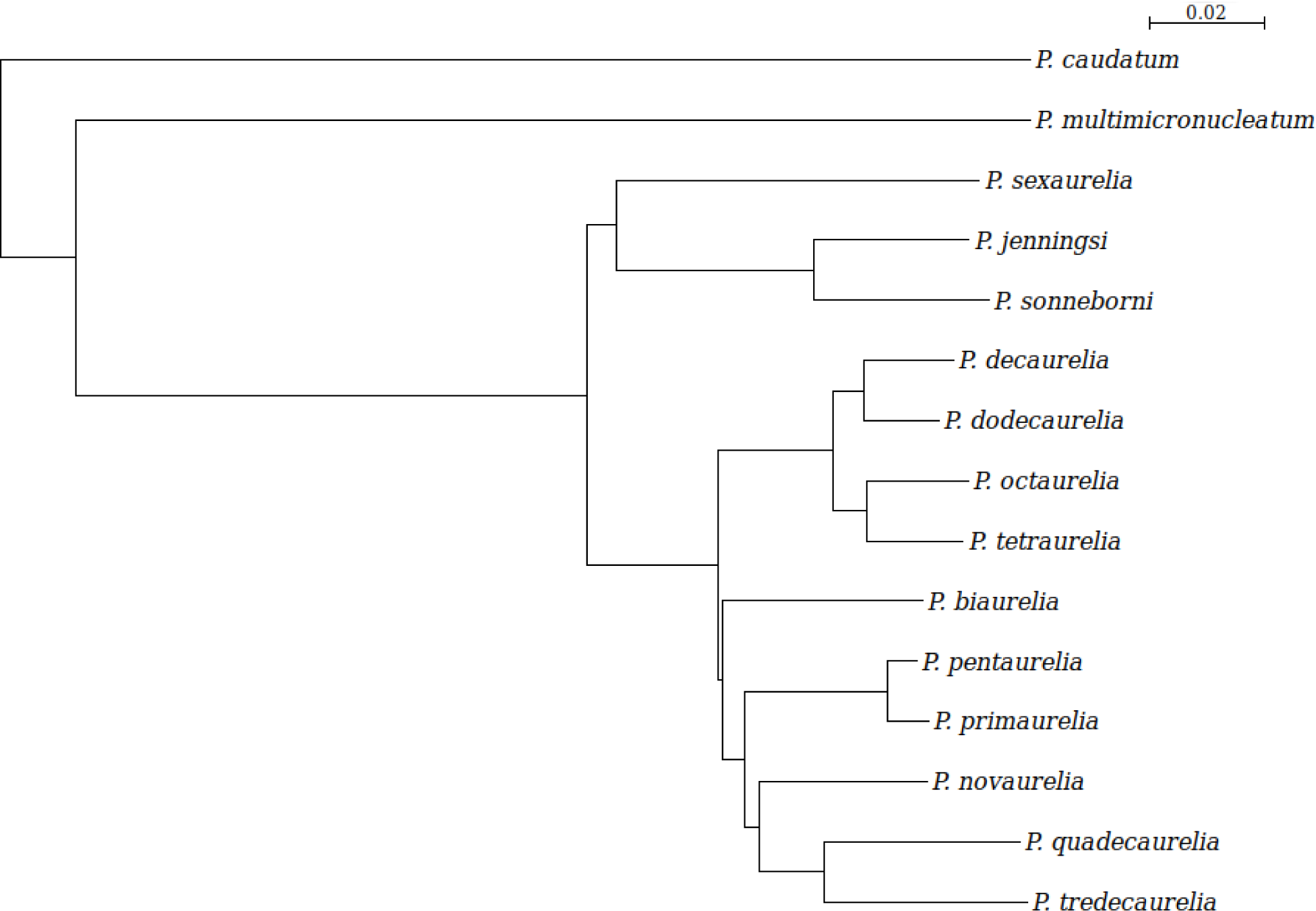
A. Phylogenetic tree of 13 species from the *P. aurelia* complex and two outgroup species. Tree based on alignment of protein-coding sequences for orthologous nuclear genes present in at least half of the *P. aurelia* species sampled (21,720 sites samples). The tree was built using the distance method implemented in seaview (Gouy, et al. 2010). Distance is amino acids substitutions.

### Timing of post-WGD gene losses

Having established the phylogenetic relationships between all 13 *P. aurelia* species sequenced, we used the patterns of gene presence/absence in the extant species to infer the timing of gene loss. Using a parsimony-based method to map the location of gene losses onto the *P. aurelia* phylogeny (Methods), we found that, following a short period of rapid gene loss immediately after the WGD, gene loss rate decreased and entered a phase of linear decline (Figure 2). This pattern is similar to what has been observed in yeast, teleost fish and plants (Scannell, et al. 2006; Inoue, et al. 2015; Ren, et al. 2018) although the exact shape of the survival curve is disputed (Inoue, et al. 2015). Using all of the ancestrally reconstructed duplicated gene survival rates, we found that an exponential decay model provided a better fit (R^2^=0.49) than a linear decay model (R^2^=0.44). However, after excluding the deepest points in the ancestral reconstruction, the exponential decay model did not outperform the linear model, suggesting a two-phase model for gene loss rate, similar to what has been reported in teleost fish (Inoue, et al. 2015). The observation that post-WGD gene-loss rate evolution follows similar trends across organisms as diverse as *Paramecium*, yeast and teleost fish suggests the possibility that the selective pressures responsible for gene retention following genome duplications are similar across the tree of eukaryotes. Although the rate at which ohnologs are lost has considerably slowed down, gene losses remain frequent in *Paramecium*. For example, we found 146 genes that have been lost in *P. primaurelia* while still being retained in two copies in the closely related sister species *P. pentaurelia*, highlighting the fact that gene loss is still an active ongoing process in *Paramecium*.

**Figure 2.**
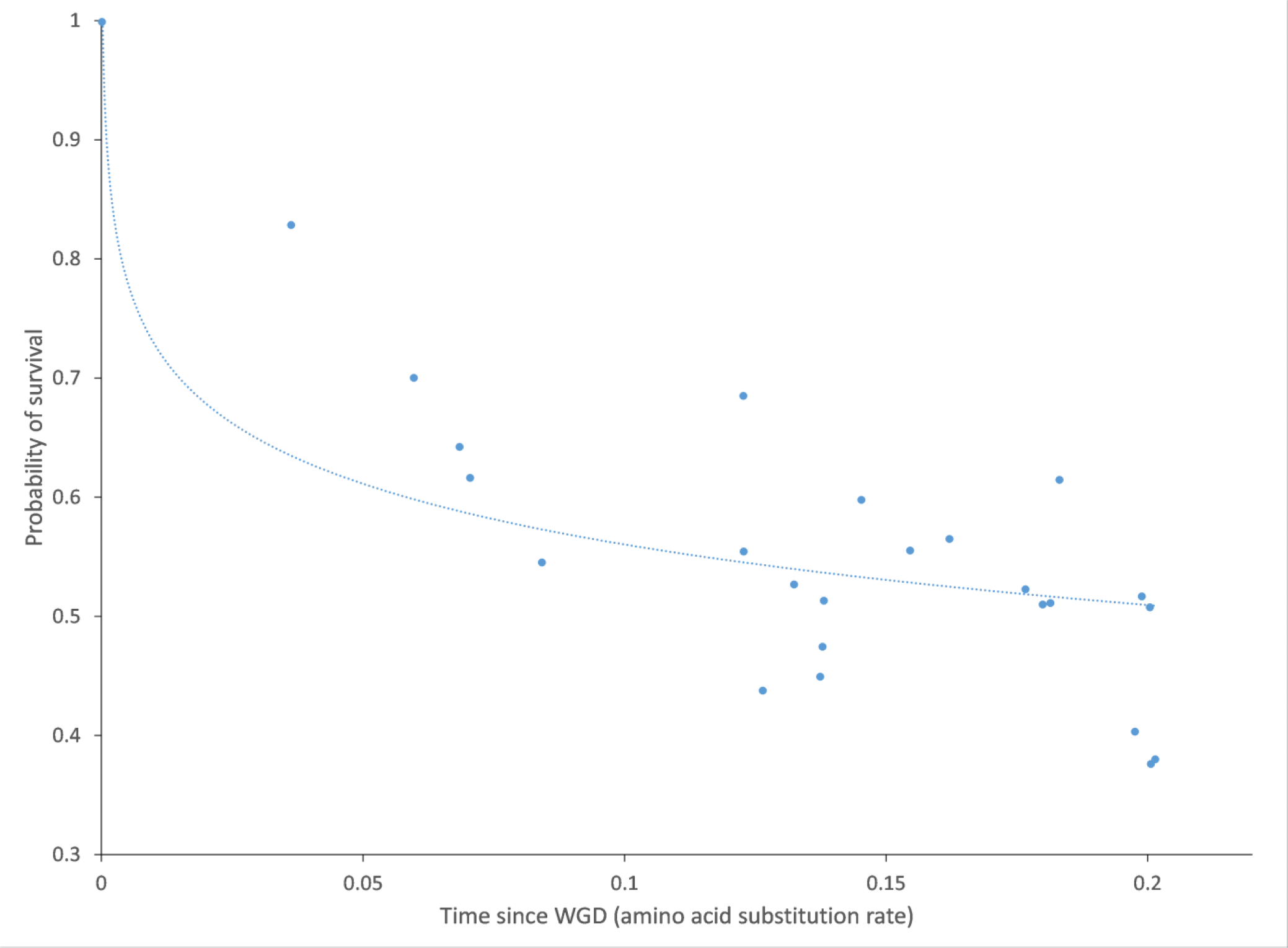
Survival curve of WGD-derived paralogs in *P. aurelia*. Ancestral retention/loss rates were estimated at each node in the tree using a parsimony-based algorithm and plotted as a function of the distance between the corresponding node and the most common ancestor of all *P. aurelia* species (which coincides with the most recent WGD (McGrath, Gout, Johri, et al. 2014)).

### Slow gene losses in *Paramecium* in comparison to other species

Having determined the general pattern of post-WGD gene loss with time, we aimed at directly comparing the strength of selective pressures responsible for ohnolog retention in different lineages. Although the amount of time elapsed since the genome duplication will be the same in all extant species, the mutation rate, generation time, and strength of selection might vary between *P. aurelia* species, resulting in different strength of selective pressures on post-WGD ohnologs evolution across these lineages. We used the average amount of synonymous substitutions between the retained ohnologs as a proxy for the amount of mutational pressure that has been faced by ohnologs since the genome duplication. Within extant *P. aurelia* species, we found a strong negative correlation between the probability of ohnolog retention and the level of sequence divergence between the remaining pairs of ohnologs (r = −0.96, p<0.01; Figure 3, red dots). This correlation remained significant when accounting for the phylogenetic non-independence of the data (r = −0.75, p = 0.003). We then performed the same analysis in a number of non-*Paramecium* lineages having experienced a genome duplication. We found that, despite a shared general trend of decreased retention with synonymous sequence divergence, the rate of post-WGD gene loss relative to synonymous divergence was lower in *Paramecium* than in other species (Figure 3). In other words, the rate of gene loss per synonymous substitution was lower in *Paramecium* than in other phylogenetic groups having experienced a genome duplication. We interpret this observation as evidence that the strength of selection opposing gene loss is stronger in *Paramecium* than in plants and vertebrates.

**Figure 3.**
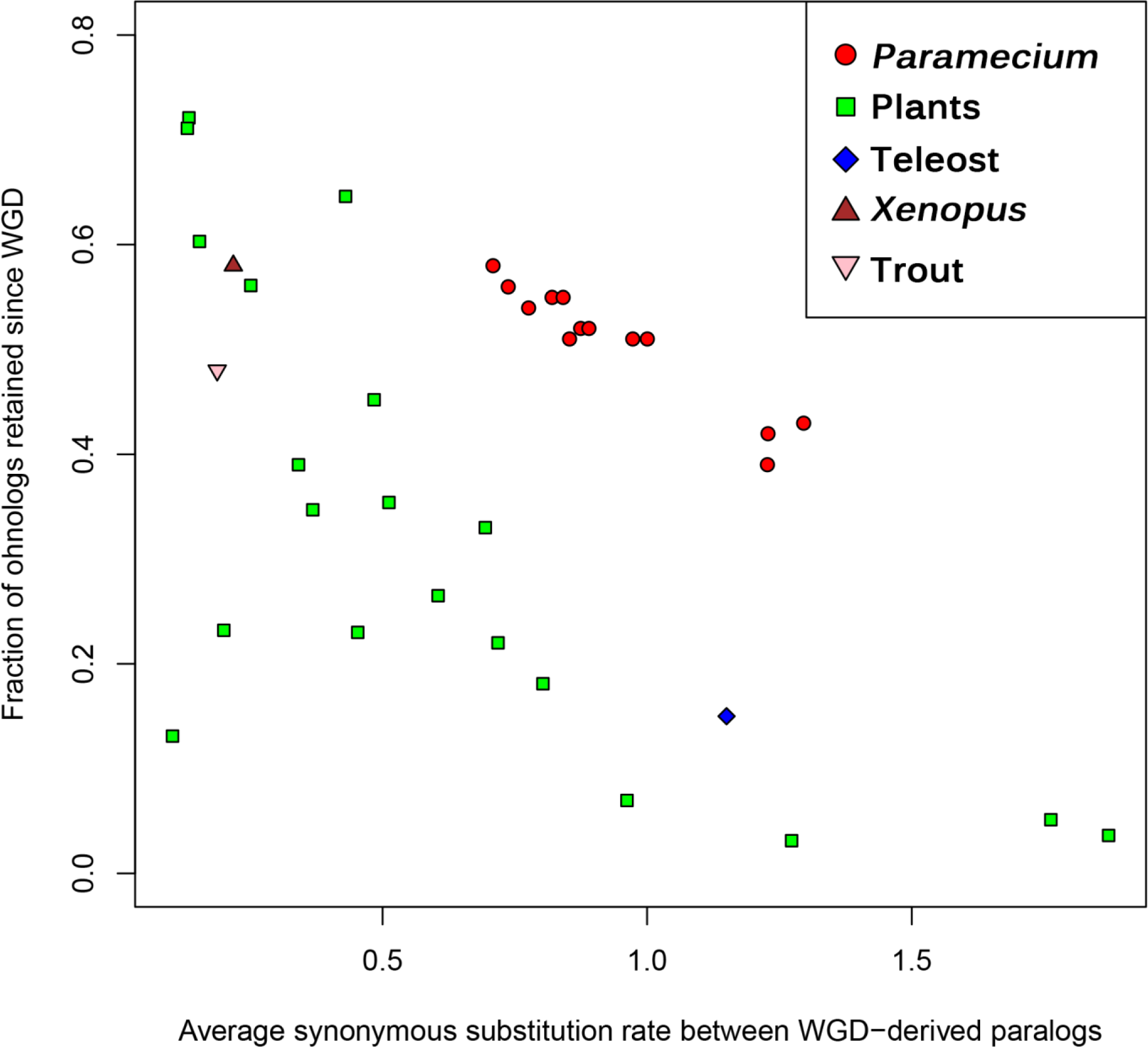
Probability of post-WGD retention as a function of sequence divergence (synonymous substitution rate) between the remaining pairs of WGD-derived paralogs in *P. aurelia* and other eukaryotes having experienced ancestral WGD.

### Selective pressures opposing gene loss

In order to understand why selection against gene loss is stronger in *Paramecium* than in other species we must first clarify the nature of the selective pressures promoting ohnolog retention. While it is difficult to pinpoint which scenario (neo/subfunctionalization, or dosage constraint) is responsible for the retention of each ohnolog pair, some general trends can be derived from genome-wide analyses. We initially reported that the probability of retention is positively correlated with the expression level of ohnologs in *Paramecium* and have interpreted this observation as evidence for stronger dosage constraints in highly expressed genes (Gout, et al. 2010; Gout and Lynch 2015). A similar trend had been reported for *S. cerevisiae* (Seoighe and Wolfe 1999), suggesting a universal role for expression level in post-WGD gene retention. We first confirmed that the increased retention rate for highly expressed genes was a universal pattern, present in all 13 *P. aurelia* species (Figure S1). With 13 *P. aurelia* species available, we were able to compute an across-species ohnolog retention rate defined as the fraction of species within the *P. aurelia* complex retaining both copies of an ohnolog pair. We used the expression level of the ortholohous gene in *P. caudatum* as a proxy for the pre-duplication expression level. As expected, we found a positive correlation between expression level and across-species retention rate (r = 0.24, p<0.001). Although this corresponds to only 6% of the variance in *P. aurelia* retention rate being explained by expression level in *P. caudatum*, the effect is consistent across all ranges of expression levels, as can be seen when genes are binned according to their expression level (Figure 4). As a result, the most highly expressed genes are twice as likely to be retained in a *P. aurelia* species than the genes with the lowest expression levels (0.81 vs 0.37 across-species retention rate for the highest and lowest bin respectively, p<0.001; Welch two sample t-test).

**Figure 4.**
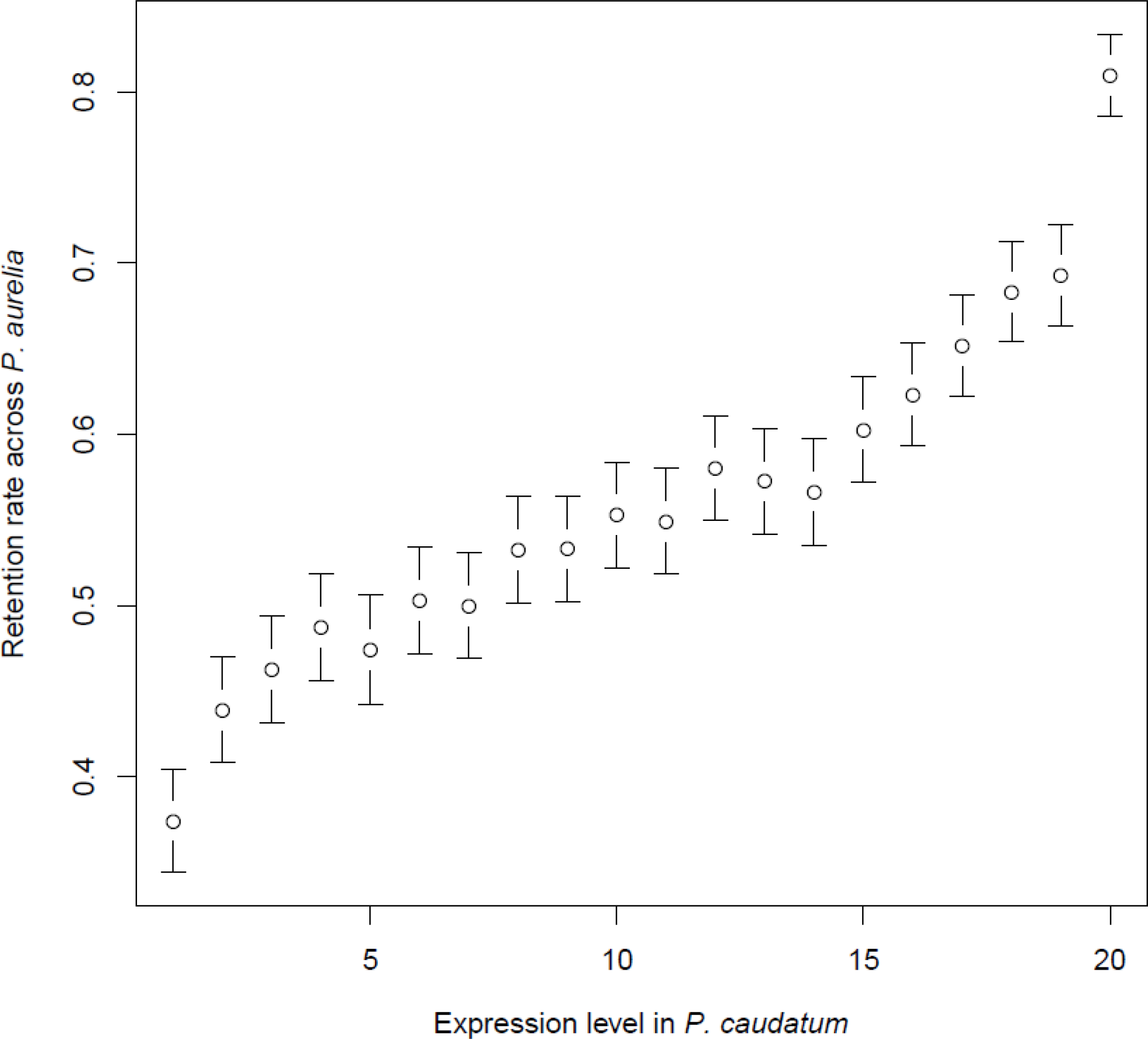
Retention rate across *P. aurelia* species as a function of orthologs expression level in *P. caudatum*. *P. caudatum* genes were classified into 20 bins of equal size according to their expression level. For each *P. caudatum* gene with an ortholog in at least on *P. aurelia* species, a retention rate was computed as the number of *P. aurelia* species where both copies have been retained divided by the number of *P. aurelia* species with at least one ortholog for this gene. Average retention rates were computed for each bin and reported on this graph alongside the 95% confidence interval.

Previous studies reported a bias in the probability of post-WGD retention for different functional categories (Seoighe and Wolfe 1999; Maere, et al. 2005; McGrath, Gout, Doak, et al. 2014; McGrath, Gout, Johri, et al. 2014; Rody, et al. 2017). We assigned Gene Ontology (GO) terms to genes in the *P. aurelia* complex and the two outgroup species using the panther pipeline (Mi, et al. 2013). We found that the retention biases per GO category were highly conserved among species, even between those that diverged very shortly after the WGD. When comparing the average retention rate for each GO category between the two groups of species that diverged the earliest (*P. sexaurelia*, *P. jeningsi*, *P. sonneborni* vs. every other species), we found a striking positive correlation (r = 0.85, p<0.01) between the two groups, indicating that the different selective pressures associated to each functional category have been preserved throughout the evolution of the *P. aurelia* complex. While the different average expression levels for each functional category explain a part of this pattern (for example genes annotated as “structural constituent of the ribosome – GO:0003735” tend to be highly expressed, and therefore are preferentially retained in two copies) we still find a number of functional categories with either significant excess or scarcity of post-WGD retention when expression level is taken into account (Table S2). One possible explanation for this pattern is that functional categories that are enriched for protein-coding genes encoding subunits of multimeric protein complexes (such as the ribosome for example) are preferentially retained due to increase dosage-balance constraints on these genes.

### Increased predetermination of paralogs’ fate

The previous observations suggest that the fate of ohnologs is at least partially predetermined at the time of duplication by their expression level and functional category. While this allows us to predict which pairs of ohnologs are most likely to rapidly lose a copy, it does not inform us about which copy, if any, is more likely to be lost. To investigate the extent of asymetrical gene loss and its evolution with time, we estimated the fraction of parallel and reciprocal gene loss at different points of the *P. aurelia* phylogeny. Parallel gene losses are cases where two species independently lose the same copy in a pair of ohnologs. Reciprocal losses are defined as two species losing a different copy in a pair of ohnologs. Gene losses that happened shortly after the genome duplication are equally distributed between reciprocal and parallel losses, as expected if both copies in a pair of ohnologous genes are equally likely to be eventually lost. However, the fraction of gene losses occupied by parallel losses increases with the distance between the genome duplication and the time of speciation between the two species considered (r = 0.4, p<0.001). In other words, one of the two genes in a pair of ohnologs becomes gradually more likely to be the one that will eventually be lost. This observation suggests that ohnologs gradually accumulate mutations that set the two copies on different trajectories, one with increased vulnerability to be eventually lost.

We previously reported that drift in expression level between ohnologs can result in a pattern such that the copy with the lowest expression is more likely to be rapidly lost (Gout and Lynch 2015). With 13 species available, we confirm that this pattern is universal across the *P. aurelia* species. Indeed, we found that, for all 13 *P. aurelia* species available, changes in expression level between two ohnologs retained in a given species increase the probability of gene loss for the orthologous pair in the closest sister species (Table S3). Additionally, gene loss is biased towards the ortholog of the copy with the lowest expression level in the sister species, a bias that becomes stronger when looking at closely related species (Table S3). For example, when looking specifically at ohnologs that have been retained in *P. decaurelia*, we find that only 3% of the orthologous pairs in *P. dodecaurelia* (the most closely related species in our dataset) have lost a copy. However, among the *P. decaurelia* ohnologs that have divergent expression level (top 5% most divergent pairs), 22% are orthologous to a *P. dodecaurelia* “pair” where one copy has been lost. This significant increase in probability of gene loss (p<0.001, χ^2^ test) is driven preferentially by the loss of the copy orthologous to the lowly expressed copy in the species that has retained both ohnologs (82% of the cases vs. 50% expected by chance, p<0.001; 1-sample proportions test with continuity correction). Therefore, it appears that divergence in gene expression between ohnologs sets the two copies on opposite trajectories for their long term survival. However, contrary to our previous prediction (Gout and Lynch 2015), we did not find any evidence for compensatory mutations increasing expression level of the remaining copy. Therefore it is possible that decreased expression level in one copy is allowed simply by reduced dosage requirements, rather than by compensatory increased expression level in the other copy.

### Interplay with segmental duplications

While *Paramecium* seems highly permissive to genome duplications, it has been noted that additional segmental duplications are rare (Aury, et al. 2006; McGrath, Gout, Doak, et al. 2014). We searched for evidence of recent segmental duplication in all *P. aurelia* genomes reported here and confirmed their extreme paucity in *P. aurelia* with a median of 28 recent segmental duplications per genome (Table S4 and Methods). This number is in sharp contrast to the thousands of gene losses that have happened since the most recent *Paramecium* WGD in each of these lineages. Despite the small number of recent segmental duplications, we were able to detect a bias for these segmental duplications to encompass genes that have already lost their ohnolog from the most recent *Paramecium* WGD. Genes that had reverted to single copy status since the WGD are on average twice as likely to be part of a subsequent recent segmental duplication as those that had maintained both WGD-derived duplicates (Table S4).

Although this observation may seem paradoxical, it is actually compatible with dosage constraints playing a major role in the fate of duplicated genes in *Paramecium*. Indeed, whole-genome duplications preserve the relative dosage between genes and might even not result in any immediate change in the amount of DNA being transcribed in *Paramecium* because of the separation between somatic and germline nuclei (McGrath, Gout, Doak, et al. 2014), a singularity shared by all ciliates. While loss of a copy would result in a dosage change following WGD, the opposite is true for segmental duplications, where it is the fixation of a new copy that would disturb the initial dosage. Therefore, if selection against dosage changes is particularly strong in *Paramecium*, as suggested by its higher rate of post-WGD gene retention compared to other species, it follows that segmental duplications should be rare. We interpret these observations as additional evidence in support of dosage sensitivity playing a major role in gene retention and duplication in *Paramecium*. Indeed, the genes that have allowed a copy to be lost following the recent WGD are also more permissive to following segmental duplications, suggesting that they are simply more tolerant to dosage changes.

## Conclusions

With this study, we now have a view of post-duplication genome evolution in 13 *Paramecium* species that share a common whole-genome duplication (WGD). All species have undergo massive gene loss since the WGD, to the point that 40 to 60% of paralogs created by the WGD (ohnologs) have lost one copy. Despite this significant variation in retention rate between species, we observed a number of strikingly similar trends in gene retention and loss across all 13 *P. aurelia* species. Most notably, highly expressed genes are systematically over-retained in two copies. Different functional categories of genes also showed consistent patterns of over- and under-retention across the entire phylogeny of *P. aurelia*. The observation that both expression level and functional category influence the probability of post-WGD retention in a way that is consistent across many species indicates that the fate of ohnologs is in part predetermined, as opposed to being dependent on the random chance of acquiring mutations that would change the function of one copy or the other. While we cannot exclude the possibility that the number of mutational targets for neo- and subfunctionalization depends on the expression level and functional category, the patterns observed here are at odds with random mutations creating new functions as the main force driving post-WGD gene retention. It should also be noted that, with the exception of genes lost very early following the genome duplication, purifying selection has been operating to maintain duplicated copies for some time before allowing gene loss. We found an average dN/dS between onhologs in *P. aurelia* species of 0.05, indicating strong purifying selection against pseudogenization operating since the WGD. Yet, many of these ohnologs have recently been lost as evidenced by the differential gene retention between closely related species, indicating that purifying selection opposed gene loss for significant amounts of time before eventually allowing gene loss. Assuming that neo- or sub-functionalization is responsible for the initial retention of a given gene, one would have to invoke a change in the strength of selection against loss of function to explain its subsequent loss. While this is not impossible (changes in the environment can alter the strength of selection on different functions for example), a more parsimonious explanation of our observation is that dosage constraints are the major driver of post-WGD gene retention pattern. Indeed, the early loss of paralogs with the lowest levels of selection for dosage constraints is expected to impact dosage requirements for their interacting partners, paving the way for future paralog gene losses. We also note that expression level is relatively fluid, with divergence in expression levels between paralogs resulting in decreased selective pressures to maintain the copy with the lowest expression level. With all these observations pointing at dosage constraints being a major contributor of post-genome duplication evolution, we interpret our finding that post-WGD gene loss relative to sequence divergence was lower in *Paramecium* than in other species with well-studied WGDs as evidence that *Paramecium* species are especially sensitive to small-scale gene dosage changes. This conclusion is corroborated by the scarcity of segmental duplications in the genome of all the *Paramecium* species studied here. In all 13 species, the few segmental duplications observed are biased towards encompassing genes that have already lost their paralog from the most recent WGD, which we interpret as additional evidence for these genes being simply more tolerant to dosage changes.

Finally, we hope that this dataset will be useful to other researchers studying whole-genome duplications and will help establish *Paramecium* as a model species for WGDs, alongside yeast.

## Materials and Methods

### Genome sequencing, assembly and annotation

Paramecium cells that had recently undergo autogamy (a self-fertilization process that create 100% homozygous individuals) were grown in up to 2 liters of Wheat Grass Powder medium (Pines International) medium before being starved and harvested. *Paramecium* cells were separated from the remaining food bacteria by filtration on a 10 µm Nitex membrane. Macronuclei were isolated away from other cellular debris by gentle lysis of the cell membrane and sucrose density separation. DNA was extracted and purified using a CTAB protocol (Doyle 1987). DNA libraries were constructed with the Illumina Nexttera DNA library preparation kit following manufacturer’s recommendations and sequencing was performed on a HiSeq 2500 machine producing 2 × 150 nt reads. Reads were trimmed for adapter sequences and quality (3′ end trimming down to Q=20) with cutadapt version 1.15 (Martin 2011). Genome assembly was performed with SPades version 3.11 (Nurk, et al. 2013) with default parameters. Final assembly was cleaned up by removing short scaffolds (less than 1 kb) and scaffolds with strong blast hits to bacterial genomes. Genome annotation was done with the EuGene pipeline (Foissac 2008) using the RNAseq data (see below) generated for each data as described in (Arnaiz, et al. 2017). The list of *Paramecium* strains used in this study is as follows. *P. primaurelia* Ir4-2, *P. biaurelia* V1-4, *P. tetraurelia* 51, *P. pentaurelia* 87, *P. sexaurelia* AZ8-4, *P. octaurelia* K8, *P. novaurelia* TE, *P. decaurelia* 223, *P. dodecaurelia* 274, *P. tredecaurelia* d13-2 (derived from 209), *P. quadecaurelia* N1A, *P. jenningsi* M, *P. sonneborni* ATCC30995, *P. multimicronucleatum* MO 3c4 and *P. caudatum* 43.

### RNAseq and expression level quantification

Paramecium cells were grown in ∼1 liter of Wheat Grass Powder medium to mid-log phase before harvesting. Cells were purified away from bacteria by filtration on a 10 µm Nitex membrane. Whole-cell RNA was isolated using TRIzol (Ambion) and the manufacturer’s suggested protocol for tissue culture cells. cDNA libraries were prepared with the Illumina TruSeq library preparation kit following the manufacturer’s suggested protocol and then sequenced with Illumina single-end 150 nt reads. RNAseq reads were mapped to each corresponding genome with Bowtie/TopHat (Langmead, et al. 2009; Kim, et al. 2013) and transcript abundance (FPKM) was computed using cufflinks (Trapnell, et al. 2010) with –multi-read-correct and –frag-bias-correct options to obtain values of FPKM for each predicted protein-coding gene. Expression level was defined for each gene as the log(FPKM+0.1), the small offset (0.1) being added to include genes with FPKM values of zero even after log-transformation.

### Orthology and paralogy relationships inference

Paralogs derived from the three successive *Paramecium* whole-genome duplications (WGDs) were annotated using the pipeline initially described in (Aury, et al. 2006). Briefly, this pipeline blast reciprocal best hits to identify large blocks of synteny derived from the most recent WGD and identify retained and lost duplicates within these blocks. Ancestral (pre-WGD) genome reconstruction is then performed by fusion of the paralogous blocks with the following criteria: if both paralogs are still retained, one copy is randomly chosen to be incorporated in the ancestral genome, if one copy has been lost, the remaining copy is included at the ancestral locus. The process is then repeated with the ancestrally reconstructed genome for more ancient genome duplications.

Orthologs were assigned using a combination of PoFF (Lechner, et al. 2014) and in-house scripts. PoFF was used to perform an initial round of orthologs prediction across all 13 *P. aurelia* species. Following this first round, an ‘orthology score’ was attributed to each pair of scaffolds linked by at least one orthologous gene pair. The score was defined as the number of genes being annotated as orthologous between the two scaffolds by PoFF. Orthology relationships were then updated with the following criteria: 1-to-2 orthology relationships where the “2” corresponds to two WGD-derived paralogs were converted to 1-to-1, selecting the gene on the scaffold with the highest orthologous score as being the ortholog. Orthology relationship with *P. caudatum* and *P. multimicronucleatum* were then inferred by selecting the genes in these two species with the highest blast hit scores to the entire *P. aurelia* orthologs family.

### Building the phylogenetic tree

Protein sequences for orthologous genes that were present in a single copy in at least half of the *P. aurelia* species were aligned to their corresponding orthologous sequences from *P. caudatum* and *P. multimicronucleatum*, using MUSCLE version 3.8 (Edgar 2004). Alignments were cleaned using gblocks (Castresana 2000) and a phylogenetic tree was build using the distance method implemented in Seaview (Gouy, et al. 2010).

### Inferring loss of gene duplicate

Branch-specific loss of gene duplicates were inferred by parsimony using ancestral reconstruction with in-house scripts. We assumed that probability of gain of duplicates is zero. Missing data was encoded as “NA” such that: ancestor(child1=“NA” and child2=“gene duplicate present”) = “gene duplicate present”; ancestor(child1=“NA” and child2=“only one duplicate present”) = “NA”; ancestor(child1=“NA” and child2 = “NA”) = “NA”. In total, 9983 gene duplicate pairs were present in the ancestor (or root) of all *P. aurelia* species. Probability of survival was obtained for every node in the phylogenetic tree (based on protein sequences) as: 1.0 – (Number of duplicates present in root – Number of duplicates present at the node) / Number of duplicates present in root.

### Finding segmental duplications

We started the search for recent segmental duplication in each species with a BLAST (Altschul, et al. 1990) search of a database containing all protein-coding genes against itself. After removing self-hits, we selected pairs of reciprocal best blast hits and removed the pairs that were already annotated as being WGD-derived paralogs. We then removed hits that were not inside a paralogon (a bloc of WGD-related genes with preserved synteny) to avoid the possibility of “contamination” with WGD-related paralogs that would have been missed by the initial annotation because of subsequent gene relocation. Finally, we computed the rate of synonymous substitution for each remaining pair of genes and retained only those with a synonymous substitution below 1.0. *P. sonneborni* was excluded from this analysis because of the presence of micronucleus-derived sequences in the genome assembly mimicking recent paralogs.

## Acknowledgments

This work was supported by National Science Foundation grant EF-0328516-A006 to M.L. Additional support for genome sequencing was provided to S.D. by France Genomique.

---

**Table S1.** Assembly statistics and average post-recent-WGD retention rate in all *Paramecium* species.

| Species | Assembly Size (Mbp) | #scaffolds | #protein-coding genes | %WGD1 paralogs retained |
| --- | --- | --- | --- | --- |
| *P. biaurelia* | 76.98 | 2362 | 40261 | 54.07 |
| *P. decaurelia* | 71.91 | 1076 | 40810 | 52.08 |
| *P. dodecaurelia* | 71.63 | 1147 | 41085 | 51.44 |
| *P. jeningsi* | 65.35 | 984 | 37098 | 58.3 |
| *P. novaurelia* | 64.79 | 1877 | 35534 | 54.87 |
| *P. octaurelia* | 72.98 | 585 | 38668 | 50.98 |
| *P. pentaurelia* | 87.29 | 310 | 41676 | 55.03 |
| *P. primaurelia* | 71.02 | 677 | 34474 | 52.48 |
| *P. quadecaurelia* | 59.12 | 710 | 33793 | 42.41 |
| *P. sexaurelia* | 68.02 | 547 | 36094 | 43.03 |
| *P. sonneborni* | 98.06 | 572 | 49951 | 56.43 |
| *P. tetraurelia* | 72.10 | 697 | 40460 | 50.71 |
| *P. tredecaurelia* | 65.93 | 245 | 36179 | 39.00 |

**Table S2.** Average retention rate for different functional categories. For each functional category with at least 20 genes in *P. caudatum*, the post-WGD retention rate was computed across all *P. aurelia* species (column #3). We then randomly draw the same number of genes from the rest of the genome (*i.e.*, genes that are not in this functional category) with similar expression levels as the genes from the functional category considered. We then compute the average retention for these randomly drown genes and repeat the random drawing 100 times to obtain an average retention rate after correction for expression level (column #4). Column #5 is the standard deviation of the retention rate across 100 random drawings.

| Gene Ontology ID | Number of *P. caudatum* genes in this functional category | Average retention rate | Average retention rate after correction for expression level | standard deviation |
| --- | --- | --- | --- | --- |
| GO:0007274 | 21 | 0.234042553 | 0.489689626 | 0.146531315 |
| GO:0016799 | 25 | 0.276470588 | 0.555562415 | 0.096006509 |
| GO:0006858 | 65 | 0.369047619 | 0.482291384 | 0.069070174 |
| GO:0006486 | 119 | 0.374277457 | 0.564433859 | 0.052919056 |
| GO:0042626 | 69 | 0.388059701 | 0.539896634 | 0.068484285 |
| GO:0016779 | 56 | 0.411764706 | 0.581520186 | 0.059600064 |
| GO:0006399 | 46 | 0.420054201 | 0.60267722 | 0.081979525 |
| GO:0016836 | 39 | 0.422712934 | 0.64706596 | 0.075342789 |
| GO:0004620 | 22 | 0.435028249 | 0.655461394 | 0.110010149 |
| GO:0019221 | 31 | 0.436781609 | 0.526031346 | 0.100265484 |
| GO:0007169 | 30 | 0.438297872 | 0.533760684 | 0.083166468 |
| GO:0016854 | 41 | 0.453703704 | 0.680423558 | 0.065406964 |
| GO:0003899 | 32 | 0.453815261 | 0.599618321 | 0.081966455 |
| GO:0006778 | 21 | 0.453947368 | 0.650411281 | 0.10376394 |
| GO:0009636 | 80 | 0.463035019 | 0.592689931 | 0.054312713 |
| GO:0006732 | 76 | 0.469448584 | 0.658887558 | 0.047713445 |
| GO:0004190 | 33 | 0.472140762 | 0.582711443 | 0.077145785 |
| GO:0006633 | 23 | 0.475 | 0.642629717 | 0.099765479 |
| GO:0008237 | 69 | 0.477011494 | 0.619098633 | 0.063174934 |
| GO:0008643 | 58 | 0.479757085 | 0.542718447 | 0.061275713 |
| GO:0006631 | 91 | 0.481804949 | 0.661268939 | 0.045590289 |
| GO:0008202 | 122 | 0.485148515 | 0.610632782 | 0.04114099 |
| GO:0004521 | 40 | 0.488721805 | 0.599632353 | 0.077823547 |
| GO:0016298 | 31 | 0.489082969 | 0.622817991 | 0.082156109 |
| GO:0006635 | 29 | 0.492307692 | 0.73191848 | 0.071318879 |
| GO:0043574 | 41 | 0.495670996 | 0.56825512 | 0.061771251 |
| GO:0004383 | 25 | 0.497777778 | 0.475339155 | 0.078290712 |
| GO:0008234 | 130 | 0.502819549 | 0.614184476 | 0.038226742 |
| GO:0003779 | 39 | 0.505263158 | 0.604309285 | 0.078582758 |
| GO:0016757 | 147 | 0.505543237 | 0.576523389 | 0.043393465 |
| GO:0003724 | 65 | 0.506896552 | 0.572213978 | 0.058812837 |
| GO:0016831 | 27 | 0.510067114 | 0.675515233 | 0.114128595 |
| GO:0005245 | 31 | 0.518691589 | 0.436756461 | 0.099568941 |
| GO:0005248 | 31 | 0.518691589 | 0.439318958 | 0.102508875 |
| GO:0019227 | 31 | 0.518691589 | 0.456333373 | 0.091858368 |
| GO:0004016 | 37 | 0.519637462 | 0.50148415 | 0.067961118 |
| GO:0006865 | 28 | 0.520408163 | 0.566520368 | 0.073991343 |
| GO:0008203 | 78 | 0.52265861 | 0.590909091 | 0.053399985 |
| GO:0008233 | 348 | 0.523140213 | 0.619757664 | 0.024601674 |
| GO:0008415 | 81 | 0.523972603 | 0.632519077 | 0.060206648 |
| GO:0051179 | 161 | 0.526264591 | 0.58872066 | 0.045959298 |
| GO:0005216 | 291 | 0.528110599 | 0.506636487 | 0.026677846 |
| GO:0006403 | 156 | 0.529292929 | 0.582263729 | 0.045357989 |
| GO:0005244 | 205 | 0.529553679 | 0.483209142 | 0.03128873 |
| GO:0005261 | 205 | 0.529553679 | 0.481737311 | 0.030961332 |
| GO:0016853 | 150 | 0.530883503 | 0.658453609 | 0.034698861 |
| GO:0022857 | 677 | 0.531031881 | 0.549919615 | 0.01813288 |
| GO:0005249 | 174 | 0.531163435 | 0.486572911 | 0.034521274 |
| GO:0015276 | 183 | 0.531628533 | 0.501379906 | 0.035526436 |
| GO:0016829 | 136 | 0.531858407 | 0.618611173 | 0.032537289 |
| GO:0005215 | 685 | 0.532465116 | 0.550289449 | 0.019074052 |
| GO:0004812 | 26 | 0.532967033 | 0.660018295 | 0.090349375 |
| GO:0006796 | 393 | 0.533524904 | 0.574047245 | 0.02195181 |
| GO:0015171 | 74 | 0.536776213 | 0.61199441 | 0.052106555 |
| GO:0043855 | 158 | 0.537827715 | 0.4820376 | 0.04044217 |
| GO:0006508 | 645 | 0.539671065 | 0.611549099 | 0.017638788 |
| GO:0007292 | 22 | 0.542372881 | 0.618230519 | 0.115993382 |
| GO:0007269 | 137 | 0.542801556 | 0.531304038 | 0.042007092 |
| GO:0008236 | 71 | 0.543010753 | 0.618707037 | 0.049347562 |
| GO:0016866 | 21 | 0.543046358 | 0.667572599 | 0.103376632 |
| GO:0006874 | 62 | 0.543429844 | 0.538189099 | 0.065068905 |
| GO:0008324 | 345 | 0.545021962 | 0.522376111 | 0.024849128 |
| GO:0006898 | 40 | 0.548295455 | 0.618387968 | 0.060184976 |
| GO:0004872 | 300 | 0.548682703 | 0.532608597 | 0.027427164 |
| GO:0032568 | 27 | 0.548717949 | 0.634531633 | 0.096569335 |
| GO:0006629 | 528 | 0.549334557 | 0.615724668 | 0.019473355 |
| GO:0007601 | 91 | 0.553524804 | 0.577415744 | 0.043486918 |
| GO:0006766 | 48 | 0.556430446 | 0.652849741 | 0.070247083 |
| GO:0006869 | 158 | 0.557198444 | 0.596669665 | 0.036893674 |
| GO:0016787 | 1487 | 0.558446889 | 0.597286431 | 0.012039042 |
| GO:0006310 | 36 | 0.558823529 | 0.543663358 | 0.074226274 |
| GO:0016874 | 472 | 0.559277738 | 0.591887846 | 0.021821233 |
| GO:0009110 | 40 | 0.561128527 | 0.678744995 | 0.069059856 |
| GO:0008168 | 87 | 0.561983471 | 0.594578527 | 0.042082156 |
| GO:0004518 | 115 | 0.562439497 | 0.596790342 | 0.041963497 |
| GO:0042592 | 65 | 0.5639413 | 0.554342432 | 0.065854307 |
| GO:0016407 | 49 | 0.564516129 | 0.639362059 | 0.060295687 |
| GO:0006418 | 31 | 0.565957447 | 0.628814464 | 0.101693925 |
| GO:0009566 | 32 | 0.56684492 | 0.536934576 | 0.083614356 |
| GO:0007249 | 32 | 0.567307692 | 0.593999698 | 0.104045253 |
| GO:0019882 | 38 | 0.567867036 | 0.581906808 | 0.070452149 |
| GO:0006917 | 105 | 0.569174757 | 0.561401326 | 0.045071878 |
| GO:0006811 | 549 | 0.569762123 | 0.554337419 | 0.020269084 |
| GO:0006968 | 71 | 0.570605187 | 0.596593722 | 0.051513815 |
| GO:0019207 | 354 | 0.570843291 | 0.564871232 | 0.02617709 |
| GO:0016491 | 373 | 0.571193686 | 0.671011286 | 0.020019427 |
| GO:0003924 | 114 | 0.571862348 | 0.595268636 | 0.04031664 |
| GO:0006812 | 534 | 0.573735955 | 0.553777185 | 0.017962939 |
| GO:0004842 | 253 | 0.574539363 | 0.57580668 | 0.028781897 |
| GO:0004386 | 105 | 0.574795082 | 0.575130825 | 0.041579722 |
| GO:0007267 | 314 | 0.575314657 | 0.548079255 | 0.025795913 |
| GO:0007268 | 306 | 0.577429984 | 0.53979004 | 0.027135871 |
| GO:0003746 | 44 | 0.578313253 | 0.666666667 | 0.070961136 |
| GO:0006790 | 36 | 0.579268293 | 0.638068311 | 0.068632449 |
| GO:0019210 | 187 | 0.58137931 | 0.590843215 | 0.035479906 |
| GO:0006807 | 77 | 0.584664537 | 0.649446961 | 0.050325494 |
| GO:0006644 | 178 | 0.585334199 | 0.585929365 | 0.037355682 |
| GO:0003824 | 4369 | 0.585671807 | 0.5935817 | 0.005863404 |
| GO:0009187 | 60 | 0.586715867 | 0.539175111 | 0.06083267 |
| GO:0006401 | 63 | 0.590032154 | 0.604843074 | 0.053107448 |
| GO:0006457 | 145 | 0.594240838 | 0.653879922 | 0.03628301 |
| GO:0006810 | 1611 | 0.59431988 | 0.568200769 | 0.010149306 |
| GO:0007166 | 443 | 0.596117552 | 0.572894671 | 0.019908474 |
| GO:0007186 | 295 | 0.597446489 | 0.564506516 | 0.023841852 |
| GO:0005484 | 35 | 0.597701149 | 0.608429459 | 0.080512877 |
| GO:0031201 | 35 | 0.597701149 | 0.602647715 | 0.087591898 |
| GO:0051169 | 74 | 0.597765363 | 0.625621272 | 0.05914454 |
| GO:0002504 | 31 | 0.597785978 | 0.527007907 | 0.08453823 |
| GO:0006817 | 63 | 0.598058252 | 0.619281678 | 0.056993188 |
| GO:0007155 | 52 | 0.598108747 | 0.581744745 | 0.064756165 |
| GO:0008152 | 5215 | 0.600642628 | 0.595770711 | 0.005223187 |
| GO:0003743 | 96 | 0.602739726 | 0.64547651 | 0.04478873 |
| GO:0006519 | 330 | 0.604129583 | 0.61825514 | 0.025377792 |
| GO:0006520 | 330 | 0.604129583 | 0.616299452 | 0.024928958 |
| GO:0044238 | 5024 | 0.604410336 | 0.593308509 | 0.004944078 |
| GO:0006464 | 1738 | 0.604983459 | 0.564844082 | 0.011489657 |
| GO:0006281 | 194 | 0.605921388 | 0.550584117 | 0.030666435 |
| GO:0007600 | 206 | 0.606472492 | 0.543615465 | 0.03394977 |
| GO:0003712 | 162 | 0.606725146 | 0.563643865 | 0.035959942 |
| GO:0006887 | 138 | 0.607808341 | 0.55866375 | 0.034633678 |
| GO:0050877 | 630 | 0.607819669 | 0.554466472 | 0.016475023 |
| GO:0009063 | 64 | 0.607936508 | 0.672296024 | 0.043966377 |
| GO:0022904 | 132 | 0.610287707 | 0.668519305 | 0.035544637 |
| GO:0016788 | 585 | 0.611342786 | 0.577465828 | 0.02002226 |
| GO:0016740 | 1928 | 0.61338623 | 0.567296534 | 0.010076513 |
| GO:0043234 | 66 | 0.613537118 | 0.588489516 | 0.063894711 |
| GO:0005975 | 498 | 0.615015591 | 0.612259526 | 0.021085555 |
| GO:0019538 | 2943 | 0.615181213 | 0.590044959 | 0.007519785 |
| GO:0043226 | 72 | 0.616260163 | 0.635805719 | 0.049196672 |
| GO:0007154 | 2296 | 0.616783363 | 0.554679521 | 0.008853733 |
| GO:0003008 | 712 | 0.6167915 | 0.549273085 | 0.015039702 |
| GO:0008092 | 260 | 0.617073171 | 0.565002659 | 0.029606431 |
| GO:0009950 | 47 | 0.617210682 | 0.663410984 | 0.062360064 |
| GO:0006260 | 145 | 0.618650493 | 0.581797669 | 0.041344786 |
| GO:0007165 | 2188 | 0.622179628 | 0.558366312 | 0.009927466 |
| GO:0015931 | 83 | 0.622252747 | 0.605669153 | 0.048499006 |
| GO:0008081 | 37 | 0.622807018 | 0.568220644 | 0.072849833 |
| GO:0008652 | 203 | 0.623033354 | 0.605490127 | 0.030499566 |
| GO:0003887 | 28 | 0.623616236 | 0.579070837 | 0.078586346 |
| GO:0007399 | 557 | 0.62479945 | 0.569958372 | 0.019260716 |
| GO:0045182 | 122 | 0.625137326 | 0.649301998 | 0.043009541 |
| GO:0005515 | 1428 | 0.625382263 | 0.57523976 | 0.011315686 |
| GO:0031202 | 278 | 0.625566636 | 0.609918846 | 0.027636731 |
| GO:0002376 | 1019 | 0.625942135 | 0.590968126 | 0.014635099 |
| GO:0007398 | 560 | 0.626508768 | 0.56999193 | 0.020515783 |
| GO:0000398 | 344 | 0.626629095 | 0.602907288 | 0.022438555 |
| GO:0005739 | 64 | 0.627383016 | 0.644679844 | 0.04763292 |
| GO:0001501 | 72 | 0.628571429 | 0.550598911 | 0.057162095 |
| GO:0048731 | 667 | 0.629643742 | 0.569282105 | 0.016780603 |
| GO:0009987 | 3209 | 0.629867207 | 0.553053067 | 0.008205547 |
| GO:0005743 | 58 | 0.630057803 | 0.643071858 | 0.056957086 |
| GO:0006886 | 833 | 0.631972318 | 0.574894493 | 0.01568653 |
| GO:0015031 | 833 | 0.631972318 | 0.579204231 | 0.01640927 |
| GO:0006259 | 329 | 0.632155145 | 0.573486694 | 0.025416888 |
| GO:0016301 | 1416 | 0.633410474 | 0.553227775 | 0.012313849 |
| GO:0006139 | 1835 | 0.634624116 | 0.587929306 | 0.009768593 |
| GO:0006091 | 162 | 0.634887006 | 0.684702737 | 0.034744346 |
| GO:0006928 | 238 | 0.635476464 | 0.570985452 | 0.030540153 |
| GO:0003729 | 310 | 0.636867089 | 0.598562152 | 0.027403299 |
| GO:0050896 | 780 | 0.637687897 | 0.586646382 | 0.015916131 |
| GO:0005085 | 59 | 0.637964775 | 0.541510025 | 0.069671912 |
| GO:0030234 | 739 | 0.637994554 | 0.564430286 | 0.015456359 |
| GO:0006521 | 23 | 0.63800905 | 0.586640862 | 0.097560477 |
| GO:0016209 | 30 | 0.63800905 | 0.639516916 | 0.080056556 |
| GO:0006468 | 1366 | 0.638586001 | 0.55240614 | 0.013950664 |
| GO:0003723 | 413 | 0.638662546 | 0.607727674 | 0.023955671 |
| GO:0008017 | 218 | 0.639439907 | 0.548167785 | 0.033573039 |
| GO:0004672 | 1097 | 0.641737971 | 0.551450737 | 0.012607221 |
| GO:0006350 | 750 | 0.642010264 | 0.585216082 | 0.016311325 |
| GO:0006366 | 748 | 0.642010264 | 0.585455612 | 0.015931733 |
| GO:0009790 | 107 | 0.643312102 | 0.620879244 | 0.042445539 |
| GO:0006936 | 345 | 0.643379663 | 0.539140067 | 0.025508571 |
| GO:0005488 | 2914 | 0.643445487 | 0.586063913 | 0.007077006 |
| GO:0005996 | 189 | 0.643925793 | 0.60367787 | 0.033296478 |
| GO:0003678 | 55 | 0.644 | 0.572272561 | 0.062714033 |
| GO:0007059 | 260 | 0.644465291 | 0.568690153 | 0.029326777 |
| GO:0006915 | 403 | 0.646470956 | 0.60875029 | 0.023826666 |
| GO:0003676 | 1677 | 0.647835769 | 0.60293455 | 0.011058052 |
| GO:0032502 | 1106 | 0.648040224 | 0.561716062 | 0.013519741 |
| GO:0008135 | 132 | 0.648330059 | 0.648874618 | 0.039457361 |
| GO:0016192 | 575 | 0.648353828 | 0.569967866 | 0.017890797 |
| GO:0007389 | 91 | 0.65 | 0.613883425 | 0.053042646 |
| GO:0004857 | 270 | 0.650512582 | 0.588360979 | 0.027872006 |
| GO:0006605 | 80 | 0.651669086 | 0.594734251 | 0.047966544 |
| GO:0006897 | 229 | 0.652881041 | 0.572441049 | 0.025374002 |
| GO:0003677 | 687 | 0.653489235 | 0.588190529 | 0.017724067 |
| GO:0007254 | 82 | 0.654371585 | 0.597094374 | 0.056518742 |
| GO:0007242 | 1223 | 0.655252692 | 0.571797339 | 0.013283839 |
| GO:0005634 | 21 | 0.655629139 | 0.641556633 | 0.109531348 |
| GO:0016070 | 661 | 0.655755793 | 0.595499431 | 0.018422415 |
| GO:0019722 | 458 | 0.656634747 | 0.565104404 | 0.023014894 |
| GO:0006397 | 414 | 0.658679135 | 0.606653083 | 0.019273936 |
| GO:0006950 | 605 | 0.658863545 | 0.582334583 | 0.01849788 |
| GO:0005886 | 102 | 0.65902965 | 0.484724296 | 0.046112135 |
| GO:0000003 | 272 | 0.659881255 | 0.574384954 | 0.029189579 |
| GO:0042116 | 41 | 0.663120567 | 0.635033595 | 0.078680819 |
| GO:0019208 | 44 | 0.663529412 | 0.583530722 | 0.066249604 |
| GO:0007276 | 269 | 0.663632423 | 0.567589609 | 0.024089349 |
| GO:0043066 | 200 | 0.66383257 | 0.615260605 | 0.030765342 |
| GO:0005102 | 167 | 0.664217487 | 0.625177349 | 0.031282425 |
| GO:0005083 | 165 | 0.66572836 | 0.550481345 | 0.034076465 |
| GO:0004715 | 56 | 0.666666667 | 0.580129178 | 0.061943153 |
| GO:0005874 | 473 | 0.666666667 | 0.552551454 | 0.02115745 |
| GO:0030528 | 664 | 0.667376161 | 0.569787562 | 0.017974191 |
| GO:0005516 | 296 | 0.667892157 | 0.592190768 | 0.026051485 |
| GO:0030879 | 60 | 0.668449198 | 0.545533465 | 0.060299602 |
| GO:0005509 | 368 | 0.670177749 | 0.6006682 | 0.022485066 |
| GO:0003756 | 28 | 0.671641791 | 0.625221997 | 0.086808597 |
| GO:0005737 | 108 | 0.671693735 | 0.652497429 | 0.041111563 |
| GO:0003700 | 576 | 0.672058498 | 0.580007643 | 0.020033713 |
| GO:0003697 | 46 | 0.672605791 | 0.66015194 | 0.053275155 |
| GO:0005622 | 836 | 0.672727273 | 0.573408739 | 0.015392888 |
| GO:0009653 | 575 | 0.672905332 | 0.550898962 | 0.018531763 |
| GO:0032989 | 575 | 0.672905332 | 0.549167959 | 0.019252309 |
| GO:0016043 | 723 | 0.673095945 | 0.561060783 | 0.016167764 |
| GO:0006096 | 86 | 0.673563218 | 0.615155295 | 0.041819695 |
| GO:0005200 | 606 | 0.675084175 | 0.56576527 | 0.017577759 |
| GO:0005856 | 606 | 0.675084175 | 0.562421312 | 0.020514749 |
| GO:0006996 | 150 | 0.675675676 | 0.592395598 | 0.032880664 |
| GO:0015629 | 77 | 0.676748582 | 0.600207559 | 0.053447527 |
| GO:0006325 | 137 | 0.678414097 | 0.592152793 | 0.037716483 |
| GO:0007498 | 230 | 0.682310469 | 0.568973715 | 0.032128962 |
| GO:0006119 | 33 | 0.682432432 | 0.658503604 | 0.077379535 |
| GO:0006357 | 433 | 0.683658171 | 0.593044021 | 0.022965885 |
| GO:0005976 | 121 | 0.684322034 | 0.6028503 | 0.040438251 |
| GO:0006206 | 45 | 0.684863524 | 0.6276473 | 0.066472265 |
| GO:0006094 | 63 | 0.687022901 | 0.564642647 | 0.055201761 |
| GO:0016079 | 23 | 0.687782805 | 0.599492386 | 0.098472215 |
| GO:0003688 | 24 | 0.688 | 0.627093588 | 0.081445227 |
| GO:0007049 | 1439 | 0.689782217 | 0.57040146 | 0.010824217 |
| GO:0003682 | 110 | 0.690402477 | 0.587713001 | 0.045959412 |
| GO:0006367 | 24 | 0.692307692 | 0.52697699 | 0.112649521 |
| GO:0003777 | 103 | 0.692961165 | 0.560720653 | 0.051186746 |
| GO:0000165 | 338 | 0.693843031 | 0.588014774 | 0.029385626 |
| GO:0007126 | 276 | 0.696105982 | 0.574456573 | 0.027446769 |
| GO:0007283 | 179 | 0.697766097 | 0.559780531 | 0.031268274 |
| GO:0005544 | 106 | 0.698765432 | 0.563975915 | 0.045044897 |
| GO:0007379 | 43 | 0.704607046 | 0.586684519 | 0.071930346 |
| GO:0007067 | 587 | 0.70764526 | 0.573984693 | 0.019575286 |
| GO:0006378 | 39 | 0.710743802 | 0.640230117 | 0.071484456 |
| GO:0008143 | 39 | 0.710743802 | 0.641022723 | 0.0655932 |
| GO:0031124 | 41 | 0.710743802 | 0.63981978 | 0.067402329 |
| GO:0016791 | 198 | 0.712999437 | 0.608024257 | 0.035778963 |
| GO:0003774 | 120 | 0.715351812 | 0.548914459 | 0.04606166 |
| GO:0006412 | 382 | 0.715654342 | 0.683314784 | 0.021696982 |
| GO:0005198 | 831 | 0.716534141 | 0.60439391 | 0.015125542 |
| GO:0000910 | 201 | 0.738547486 | 0.554193399 | 0.031553981 |
| GO:0006099 | 32 | 0.739776952 | 0.744715951 | 0.073005605 |
| GO:0006800 | 67 | 0.740740741 | 0.672346057 | 0.067736736 |
| GO:0016072 | 179 | 0.741491673 | 0.596550886 | 0.035791514 |
| GO:0005977 | 99 | 0.741697417 | 0.617370022 | 0.046489737 |
| GO:0004601 | 28 | 0.748663102 | 0.675695418 | 0.099283285 |
| GO:0019209 | 44 | 0.753117207 | 0.561002301 | 0.0643505 |
| GO:0004721 | 171 | 0.753716871 | 0.610050505 | 0.030731241 |
| GO:0008047 | 50 | 0.759036145 | 0.589251494 | 0.066961541 |
| GO:0006461 | 58 | 0.762781186 | 0.692280337 | 0.05406727 |
| GO:0030529 | 76 | 0.773349938 | 0.64380531 | 0.045346437 |
| GO:0019201 | 40 | 0.776978417 | 0.640616763 | 0.082829792 |
| GO:0006144 | 167 | 0.791492911 | 0.620143659 | 0.036017127 |
| GO:0006369 | 112 | 0.798029557 | 0.570815493 | 0.046496402 |
| GO:0019212 | 21 | 0.81 | 0.590968236 | 0.100055853 |
| GO:0030120 | 33 | 0.811965812 | 0.643773029 | 0.082904072 |
| GO:0006109 | 34 | 0.813688213 | 0.662244692 | 0.075235856 |
| GO:0007517 | 26 | 0.837662338 | 0.556565657 | 0.099596622 |
| GO:0003735 | 213 | 0.853010164 | 0.778815619 | 0.026152334 |
| GO:0015078 | 31 | 0.855072464 | 0.742446615 | 0.063812089 |
| GO:0019219 | 31 | 0.878661088 | 0.657460317 | 0.084951587 |

**Table S3.**
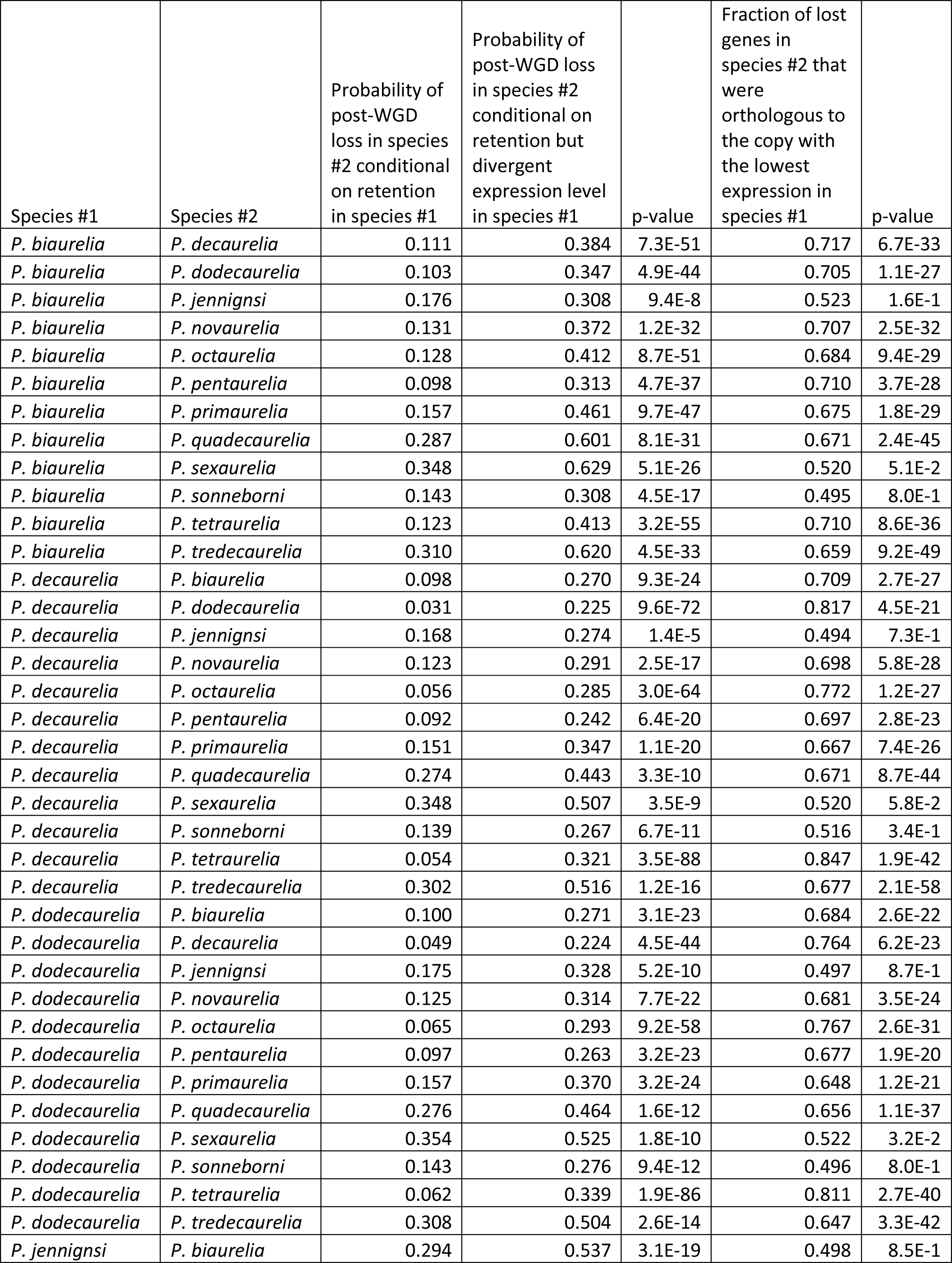
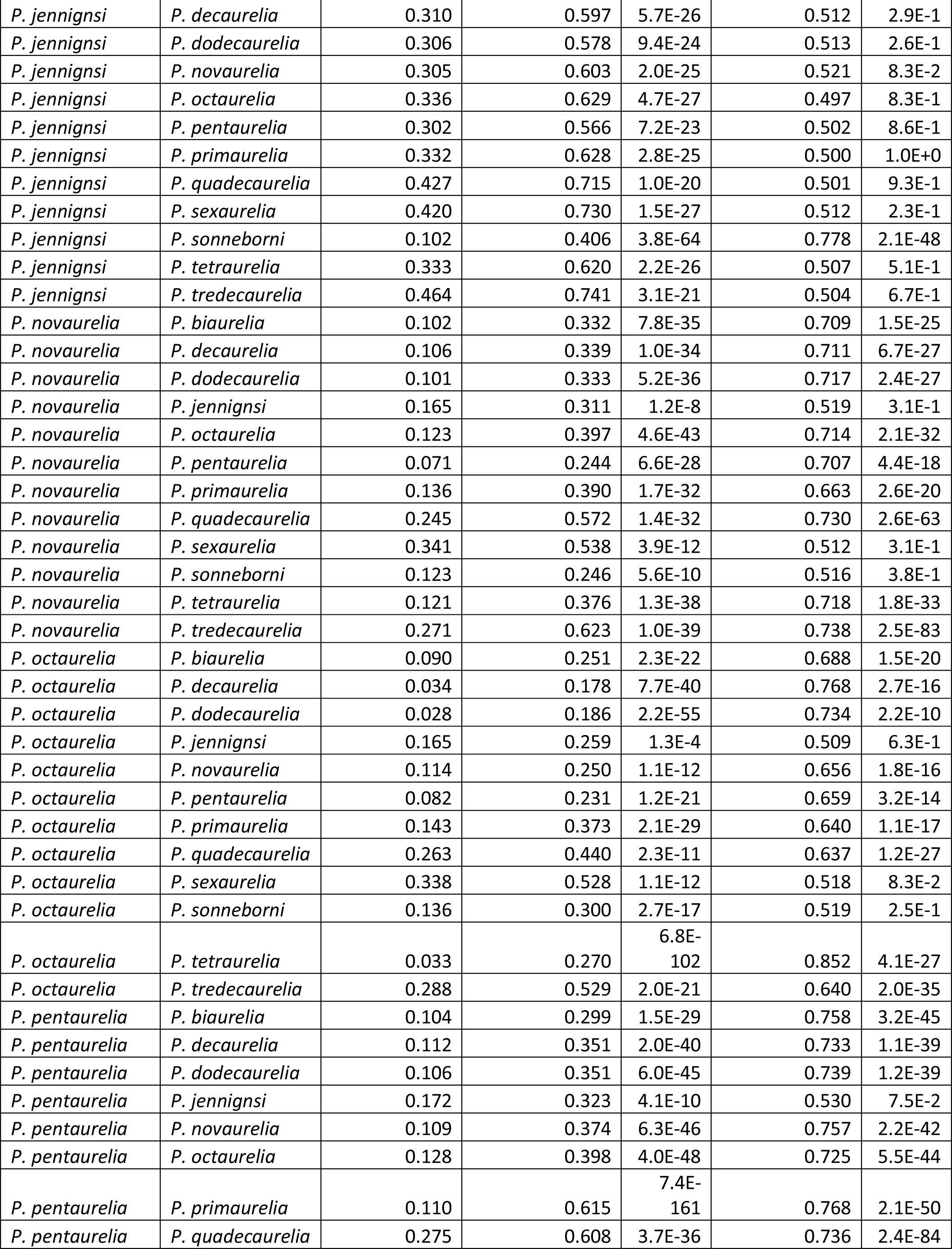
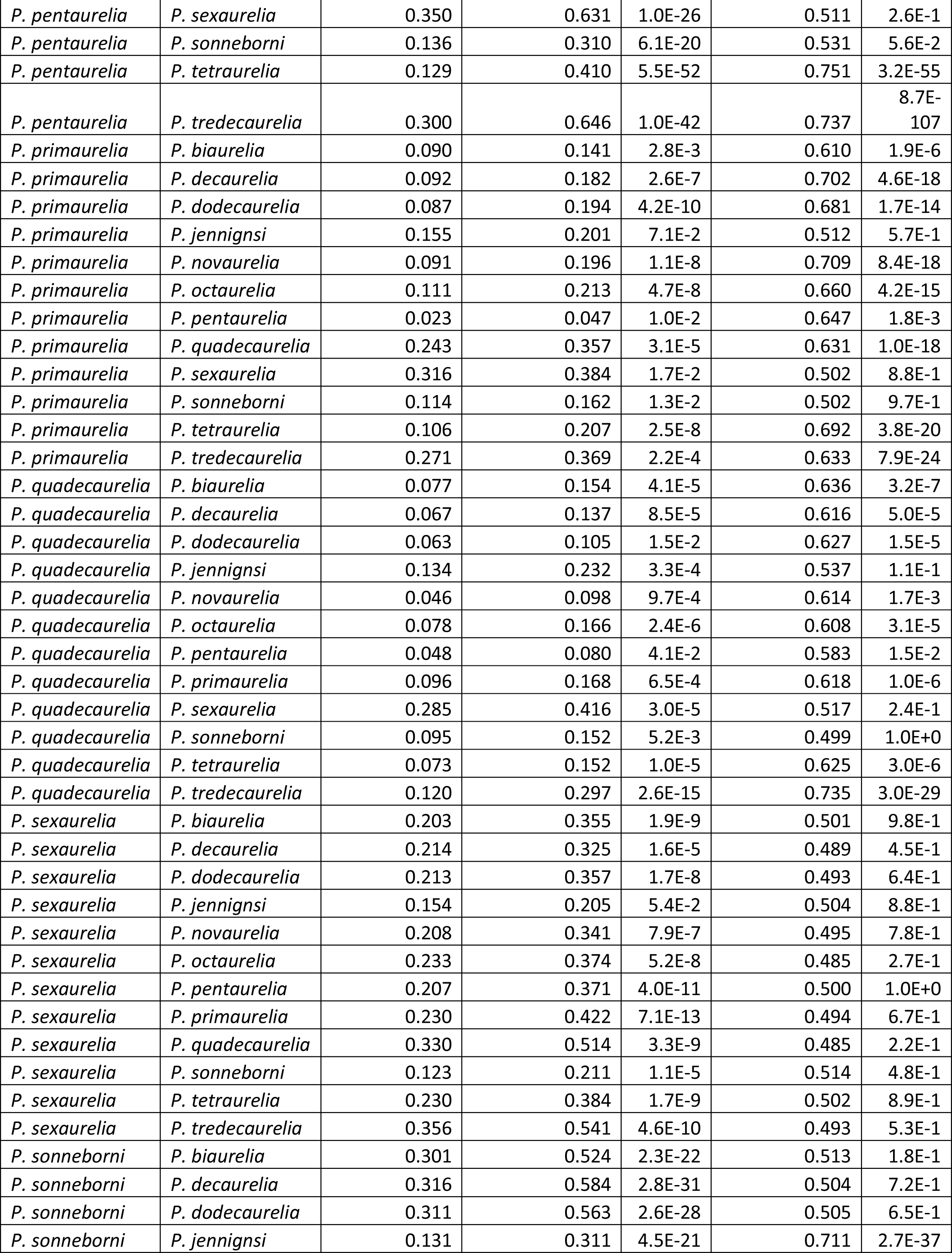
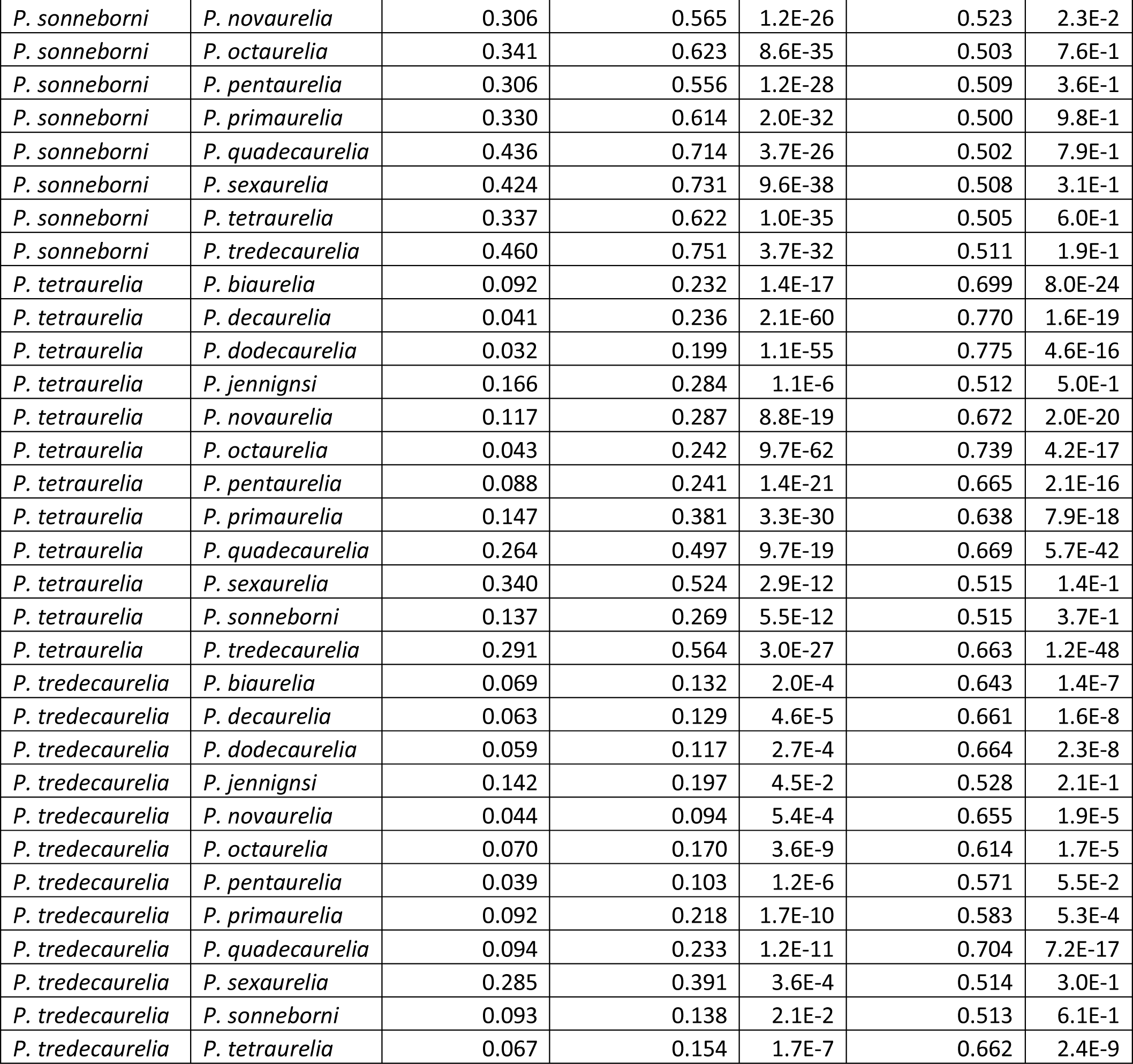
Probability of post-WGD gene loss in a given *Paramecium* as a function of retention and expression level conservation of the orthologous genes in the other *Paramecium* species.

| Species #1 | Species #2 | Probability of post-WGD loss in species #2 conditional on retention in species #1 | Probability of post-WGD loss in species #2 conditional on retention but divergent expression level in species #1 | p-value | Fraction of lost genes in species #2 that were orthologous to the copy with the lowest expression in species #1 | p-value |
| --- | --- | --- | --- | --- | --- | --- |
| *P. biaurelia* | *P. decaurelia* | 0.111 | 0.384 | 7.3E-51 | 0.717 | 6.7E-33 |
| *P. biaurelia* | *P. dodecaurelia* | 0.103 | 0.347 | 4.9E-44 | 0.705 | 1.1E-27 |
| *P. biaurelia* | *P. jennignsi* | 0.176 | 0.308 | 9.4E-8 | 0.523 | 1.6E-1 |
| *P. biaurelia* | *P. novaurelia* | 0.131 | 0.372 | 1.2E-32 | 0.707 | 2.5E-32 |
| *P. biaurelia* | *P. octaurelia* | 0.128 | 0.412 | 8.7E-51 | 0.684 | 9.4E-29 |
| *P. biaurelia* | *P. pentaurelia* | 0.098 | 0.313 | 4.7E-37 | 0.710 | 3.7E-28 |
| *P. biaurelia* | *P. primaurelia* | 0.157 | 0.461 | 9.7E-47 | 0.675 | 1.8E-29 |
| *P. biaurelia* | *P. quadecaurelia* | 0.287 | 0.601 | 8.1E-31 | 0.671 | 2.4E-45 |
| *P. biaurelia* | *P. sexaurelia* | 0.348 | 0.629 | 5.1E-26 | 0.520 | 5.1E-2 |
| *P. biaurelia* | *P. sonneborni* | 0.143 | 0.308 | 4.5E-17 | 0.495 | 8.0E-1 |
| *P. biaurelia* | *P. tetraurelia* | 0.123 | 0.413 | 3.2E-55 | 0.710 | 8.6E-36 |
| *P. biaurelia* | *P. tredecaurelia* | 0.310 | 0.620 | 4.5E-33 | 0.659 | 9.2E-49 |
| *P. decaurelia* | *P. biaurelia* | 0.098 | 0.270 | 9.3E-24 | 0.709 | 2.7E-27 |
| *P. decaurelia* | *P. dodecaurelia* | 0.031 | 0.225 | 9.6E-72 | 0.817 | 4.5E-21 |
| *P. decaurelia* | *P. jennignsi* | 0.168 | 0.274 | 1.4E-5 | 0.494 | 7.3E-1 |
| *P. decaurelia* | *P. novaurelia* | 0.123 | 0.291 | 2.5E-17 | 0.698 | 5.8E-28 |
| *P. decaurelia* | *P. octaurelia* | 0.056 | 0.285 | 3.0E-64 | 0.772 | 1.2E-27 |
| *P. decaurelia* | *P. pentaurelia* | 0.092 | 0.242 | 6.4E-20 | 0.697 | 2.8E-23 |
| *P. decaurelia* | *P. primaurelia* | 0.151 | 0.347 | 1.1E-20 | 0.667 | 7.4E-26 |
| *P. decaurelia* | *P. quadecaurelia* | 0.274 | 0.443 | 3.3E-10 | 0.671 | 8.7E-44 |
| *P. decaurelia* | *P. sexaurelia* | 0.348 | 0.507 | 3.5E-9 | 0.520 | 5.8E-2 |
| *P. decaurelia* | *P. sonneborni* | 0.139 | 0.267 | 6.7E-11 | 0.516 | 3.4E-1 |
| *P. decaurelia* | *P. tetraurelia* | 0.054 | 0.321 | 3.5E-88 | 0.847 | 1.9E-42 |
| *P. decaurelia* | *P. tredecaurelia* | 0.302 | 0.516 | 1.2E-16 | 0.677 | 2.1E-58 |
| *P. dodecaurelia* | *P. biaurelia* | 0.100 | 0.271 | 3.1E-23 | 0.684 | 2.6E-22 |
| *P. dodecaurelia* | *P. decaurelia* | 0.049 | 0.224 | 4.5E-44 | 0.764 | 6.2E-23 |
| *P. dodecaurelia* | *P. jennignsi* | 0.175 | 0.328 | 5.2E-10 | 0.497 | 8.7E-1 |
| *P. dodecaurelia* | *P. novaurelia* | 0.125 | 0.314 | 7.7E-22 | 0.681 | 3.5E-24 |
| *P. dodecaurelia* | *P. octaurelia* | 0.065 | 0.293 | 9.2E-58 | 0.767 | 2.6E-31 |
| *P. dodecaurelia* | *P. pentaurelia* | 0.097 | 0.263 | 3.2E-23 | 0.677 | 1.9E-20 |
| *P. dodecaurelia* | *P. primaurelia* | 0.157 | 0.370 | 3.2E-24 | 0.648 | 1.2E-21 |
| *P. dodecaurelia* | *P. quadecaurelia* | 0.276 | 0.464 | 1.6E-12 | 0.656 | 1.1E-37 |
| *P. dodecaurelia* | *P. sexaurelia* | 0.354 | 0.525 | 1.8E-10 | 0.522 | 3.2E-2 |
| *P. dodecaurelia* | *P. sonneborni* | 0.143 | 0.276 | 9.4E-12 | 0.496 | 8.0E-1 |
| *P. dodecaurelia* | *P. tetraurelia* | 0.062 | 0.339 | 1.9E-86 | 0.811 | 2.7E-40 |
| *P. dodecaurelia* | *P. tredecaurelia* | 0.308 | 0.504 | 2.6E-14 | 0.647 | 3.3E-42 |
| *P. jennignsi* | *P. biaurelia* | 0.294 | 0.537 | 3.1E-19 | 0.498 | 8.5E-1 |
| *P. jennignsi* | *P. decaurelia* | 0.310 | 0.597 | 5.7E-26 | 0.512 | 2.9E-1 |
| *P. jennignsi* | *P. dodecaurelia* | 0.306 | 0.578 | 9.4E-24 | 0.513 | 2.6E-1 |
| *P. jennignsi* | *P. novaurelia* | 0.305 | 0.603 | 2.0E-25 | 0.521 | 8.3E-2 |
| *P. jennignsi* | *P. octaurelia* | 0.336 | 0.629 | 4.7E-27 | 0.497 | 8.3E-1 |
| *P. jennignsi* | *P. pentaurelia* | 0.302 | 0.566 | 7.2E-23 | 0.502 | 8.6E-1 |
| *P. jennignsi* | *P. primaurelia* | 0.332 | 0.628 | 2.8E-25 | 0.500 | 1.0E+0 |
| *P. jennignsi* | *P. quadecaurelia* | 0.427 | 0.715 | 1.0E-20 | 0.501 | 9.3E-1 |
| *P. jennignsi* | *P. sexaurelia* | 0.420 | 0.730 | 1.5E-27 | 0.512 | 2.3E-1 |
| *P. jennignsi* | *P. sonneborni* | 0.102 | 0.406 | 3.8E-64 | 0.778 | 2.1E-48 |
| *P. jennignsi* | *P. tetraurelia* | 0.333 | 0.620 | 2.2E-26 | 0.507 | 5.1E-1 |
| *P. jennignsi* | *P. tredecaurelia* | 0.464 | 0.741 | 3.1E-21 | 0.504 | 6.7E-1 |
| *P. novaurelia* | *P. biaurelia* | 0.102 | 0.332 | 7.8E-35 | 0.709 | 1.5E-25 |
| *P. novaurelia* | *P. decaurelia* | 0.106 | 0.339 | 1.0E-34 | 0.711 | 6.7E-27 |
| *P. novaurelia* | *P. dodecaurelia* | 0.101 | 0.333 | 5.2E-36 | 0.717 | 2.4E-27 |
| *P. novaurelia* | *P. jennignsi* | 0.165 | 0.311 | 1.2E-8 | 0.519 | 3.1E-1 |
| *P. novaurelia* | *P. octaurelia* | 0.123 | 0.397 | 4.6E-43 | 0.714 | 2.1E-32 |
| *P. novaurelia* | *P. pentaurelia* | 0.071 | 0.244 | 6.6E-28 | 0.707 | 4.4E-18 |
| *P. novaurelia* | *P. primaurelia* | 0.136 | 0.390 | 1.7E-32 | 0.663 | 2.6E-20 |
| *P. novaurelia* | *P. quadecaurelia* | 0.245 | 0.572 | 1.4E-32 | 0.730 | 2.6E-63 |
| *P. novaurelia* | *P. sexaurelia* | 0.341 | 0.538 | 3.9E-12 | 0.512 | 3.1E-1 |
| *P. novaurelia* | *P. sonneborni* | 0.123 | 0.246 | 5.6E-10 | 0.516 | 3.8E-1 |
| *P. novaurelia* | *P. tetraurelia* | 0.121 | 0.376 | 1.3E-38 | 0.718 | 1.8E-33 |
| *P. novaurelia* | *P. tredecaurelia* | 0.271 | 0.623 | 1.0E-39 | 0.738 | 2.5E-83 |
| *P. octaurelia* | *P. biaurelia* | 0.090 | 0.251 | 2.3E-22 | 0.688 | 1.5E-20 |
| *P. octaurelia* | *P. decaurelia* | 0.034 | 0.178 | 7.7E-40 | 0.768 | 2.7E-16 |
| *P. octaurelia* | *P. dodecaurelia* | 0.028 | 0.186 | 2.2E-55 | 0.734 | 2.2E-10 |
| *P. octaurelia* | *P. jennignsi* | 0.165 | 0.259 | 1.3E-4 | 0.509 | 6.3E-1 |
| *P. octaurelia* | *P. novaurelia* | 0.114 | 0.250 | 1.1E-12 | 0.656 | 1.8E-16 |
| *P. octaurelia* | *P. pentaurelia* | 0.082 | 0.231 | 1.2E-21 | 0.659 | 3.2E-14 |
| *P. octaurelia* | *P. primaurelia* | 0.143 | 0.373 | 2.1E-29 | 0.640 | 1.1E-17 |
| *P. octaurelia* | *P. quadecaurelia* | 0.263 | 0.440 | 2.3E-11 | 0.637 | 1.2E-27 |
| *P. octaurelia* | *P. sexaurelia* | 0.338 | 0.528 | 1.1E-12 | 0.518 | 8.3E-2 |
| *P. octaurelia* | *P. sonneborni* | 0.136 | 0.300 | 2.7E-17 | 0.519 | 2.5E-1 |
| *P. octaurelia* | *P. tetraurelia* | 0.033 | 0.270 | 6.8E-102 | 0.852 | 4.1E-27 |
| *P. octaurelia* | *P. tredecaurelia* | 0.288 | 0.529 | 2.0E-21 | 0.640 | 2.0E-35 |
| *P. pentaurelia* | *P. biaurelia* | 0.104 | 0.299 | 1.5E-29 | 0.758 | 3.2E-45 |
| *P. pentaurelia* | *P. decaurelia* | 0.112 | 0.351 | 2.0E-40 | 0.733 | 1.1E-39 |
| *P. pentaurelia* | *P. dodecaurelia* | 0.106 | 0.351 | 6.0E-45 | 0.739 | 1.2E-39 |
| *P. pentaurelia* | *P. jennignsi* | 0.172 | 0.323 | 4.1E-10 | 0.530 | 7.5E-2 |
| *P. pentaurelia* | *P. novaurelia* | 0.109 | 0.374 | 6.3E-46 | 0.757 | 2.2E-42 |
| *P. pentaurelia* | *P. octaurelia* | 0.128 | 0.398 | 4.0E-48 | 0.725 | 5.5E-44 |
| *P. pentaurelia* | *P. primaurelia* | 0.110 | 0.615 | 7.4E-161 | 0.768 | 2.1E-50 |
| *P. pentaurelia* | *P. quadecaurelia* | 0.275 | 0.608 | 3.7E-36 | 0.736 | 2.4E-84 |
| *P. pentaurelia* | *P. sexaurelia* | 0.350 | 0.631 | 1.0E-26 | 0.511 | 2.6E-1 |
| *P. pentaurelia* | *P. sonneborni* | 0.136 | 0.310 | 6.1E-20 | 0.531 | 5.6E-2 |
| *P. pentaurelia* | *P. tetraurelia* | 0.129 | 0.410 | 5.5E-52 | 0.751 | 3.2E-55 |
| *P. pentaurelia* | *P. tredecaurelia* | 0.300 | 0.646 | 1.0E-42 | 0.737 | 8.7E-107 |
| *P. primaurelia* | *P. biaurelia* | 0.090 | 0.141 | 2.8E-3 | 0.610 | 1.9E-6 |
| *P. primaurelia* | *P. decaurelia* | 0.092 | 0.182 | 2.6E-7 | 0.702 | 4.6E-18 |
| *P. primaurelia* | *P. dodecaurelia* | 0.087 | 0.194 | 4.2E-10 | 0.681 | 1.7E-14 |
| *P. primaurelia* | *P. jennignsi* | 0.155 | 0.201 | 7.1E-2 | 0.512 | 5.7E-1 |
| *P. primaurelia* | *P. novaurelia* | 0.091 | 0.196 | 1.1E-8 | 0.709 | 8.4E-18 |
| *P. primaurelia* | *P. octaurelia* | 0.111 | 0.213 | 4.7E-8 | 0.660 | 4.2E-15 |
| *P. primaurelia* | *P. pentaurelia* | 0.023 | 0.047 | 1.0E-2 | 0.647 | 1.8E-3 |
| *P. primaurelia* | *P. quadecaurelia* | 0.243 | 0.357 | 3.1E-5 | 0.631 | 1.0E-18 |
| *P. primaurelia* | *P. sexaurelia* | 0.316 | 0.384 | 1.7E-2 | 0.502 | 8.8E-1 |
| *P. primaurelia* | *P. sonneborni* | 0.114 | 0.162 | 1.3E-2 | 0.502 | 9.7E-1 |
| *P. primaurelia* | *P. tetraurelia* | 0.106 | 0.207 | 2.5E-8 | 0.692 | 3.8E-20 |
| *P. primaurelia* | *P. tredecaurelia* | 0.271 | 0.369 | 2.2E-4 | 0.633 | 7.9E-24 |
| *P. quadecaurelia* | *P. biaurelia* | 0.077 | 0.154 | 4.1E-5 | 0.636 | 3.2E-7 |
| *P. quadecaurelia* | *P. decaurelia* | 0.067 | 0.137 | 8.5E-5 | 0.616 | 5.0E-5 |
| *P. quadecaurelia* | *P. dodecaurelia* | 0.063 | 0.105 | 1.5E-2 | 0.627 | 1.5E-5 |
| *P. quadecaurelia* | *P. jennignsi* | 0.134 | 0.232 | 3.3E-4 | 0.537 | 1.1E-1 |
| *P. quadecaurelia* | *P. novaurelia* | 0.046 | 0.098 | 9.7E-4 | 0.614 | 1.7E-3 |
| *P. quadecaurelia* | *P. octaurelia* | 0.078 | 0.166 | 2.4E-6 | 0.608 | 3.1E-5 |
| *P. quadecaurelia* | *P. pentaurelia* | 0.048 | 0.080 | 4.1E-2 | 0.583 | 1.5E-2 |
| *P. quadecaurelia* | *P. primaurelia* | 0.096 | 0.168 | 6.5E-4 | 0.618 | 1.0E-6 |
| *P. quadecaurelia* | *P. sexaurelia* | 0.285 | 0.416 | 3.0E-5 | 0.517 | 2.4E-1 |
| *P. quadecaurelia* | *P. sonneborni* | 0.095 | 0.152 | 5.2E-3 | 0.499 | 1.0E+0 |
| *P. quadecaurelia* | *P. tetraurelia* | 0.073 | 0.152 | 1.0E-5 | 0.625 | 3.0E-6 |
| *P. quadecaurelia* | *P. tredecaurelia* | 0.120 | 0.297 | 2.6E-15 | 0.735 | 3.0E-29 |
| *P. sexaurelia* | *P. biaurelia* | 0.203 | 0.355 | 1.9E-9 | 0.501 | 9.8E-1 |
| *P. sexaurelia* | *P. decaurelia* | 0.214 | 0.325 | 1.6E-5 | 0.489 | 4.5E-1 |
| *P. sexaurelia* | *P. dodecaurelia* | 0.213 | 0.357 | 1.7E-8 | 0.493 | 6.4E-1 |
| *P. sexaurelia* | *P. jennignsi* | 0.154 | 0.205 | 5.4E-2 | 0.504 | 8.8E-1 |
| *P. sexaurelia* | *P. novaurelia* | 0.208 | 0.341 | 7.9E-7 | 0.495 | 7.8E-1 |
| *P. sexaurelia* | *P. octaurelia* | 0.233 | 0.374 | 5.2E-8 | 0.485 | 2.7E-1 |
| *P. sexaurelia* | *P. pentaurelia* | 0.207 | 0.371 | 4.0E-11 | 0.500 | 1.0E+0 |
| *P. sexaurelia* | *P. primaurelia* | 0.230 | 0.422 | 7.1E-13 | 0.494 | 6.7E-1 |
| *P. sexaurelia* | *P. quadecaurelia* | 0.330 | 0.514 | 3.3E-9 | 0.485 | 2.2E-1 |
| *P. sexaurelia* | *P. sonneborni* | 0.123 | 0.211 | 1.1E-5 | 0.514 | 4.8E-1 |
| *P. sexaurelia* | *P. tetraurelia* | 0.230 | 0.384 | 1.7E-9 | 0.502 | 8.9E-1 |
| *P. sexaurelia* | *P. tredecaurelia* | 0.356 | 0.541 | 4.6E-10 | 0.493 | 5.3E-1 |
| *P. sonneborni* | *P. biaurelia* | 0.301 | 0.524 | 2.3E-22 | 0.513 | 1.8E-1 |
| *P. sonneborni* | *P. decaurelia* | 0.316 | 0.584 | 2.8E-31 | 0.504 | 7.2E-1 |
| *P. sonneborni* | *P. dodecaurelia* | 0.311 | 0.563 | 2.6E-28 | 0.505 | 6.5E-1 |
| *P. sonneborni* | *P. jennignsi* | 0.131 | 0.311 | 4.5E-21 | 0.711 | 2.7E-37 |
| *P. sonneborni* | *P. novaurelia* | 0.306 | 0.565 | 1.2E-26 | 0.523 | 2.3E-2 |
| *P. sonneborni* | *P. octaurelia* | 0.341 | 0.623 | 8.6E-35 | 0.503 | 7.6E-1 |
| *P. sonneborni* | *P. pentaurelia* | 0.306 | 0.556 | 1.2E-28 | 0.509 | 3.6E-1 |
| *P. sonneborni* | *P. primaurelia* | 0.330 | 0.614 | 2.0E-32 | 0.500 | 9.8E-1 |
| *P. sonneborni* | *P. quadecaurelia* | 0.436 | 0.714 | 3.7E-26 | 0.502 | 7.9E-1 |
| *P. sonneborni* | *P. sexaurelia* | 0.424 | 0.731 | 9.6E-38 | 0.508 | 3.1E-1 |
| *P. sonneborni* | *P. tetraurelia* | 0.337 | 0.622 | 1.0E-35 | 0.505 | 6.0E-1 |
| *P. sonneborni* | *P. tredecaurelia* | 0.460 | 0.751 | 3.7E-32 | 0.511 | 1.9E-1 |
| *P. tetraurelia* | *P. biaurelia* | 0.092 | 0.232 | 1.4E-17 | 0.699 | 8.0E-24 |
| *P. tetraurelia* | *P. decaurelia* | 0.041 | 0.236 | 2.1E-60 | 0.770 | 1.6E-19 |
| *P. tetraurelia* | *P. dodecaurelia* | 0.032 | 0.199 | 1.1E-55 | 0.775 | 4.6E-16 |
| *P. tetraurelia* | *P. jennignsi* | 0.166 | 0.284 | 1.1E-6 | 0.512 | 5.0E-1 |
| *P. tetraurelia* | *P. novaurelia* | 0.117 | 0.287 | 8.8E-19 | 0.672 | 2.0E-20 |
| *P. tetraurelia* | *P. octaurelia* | 0.043 | 0.242 | 9.7E-62 | 0.739 | 4.2E-17 |
| *P. tetraurelia* | *P. pentaurelia* | 0.088 | 0.241 | 1.4E-21 | 0.665 | 2.1E-16 |
| *P. tetraurelia* | *P. primaurelia* | 0.147 | 0.381 | 3.3E-30 | 0.638 | 7.9E-18 |
| *P. tetraurelia* | *P. quadecaurelia* | 0.264 | 0.497 | 9.7E-19 | 0.669 | 5.7E-42 |
| *P. tetraurelia* | *P. sexaurelia* | 0.340 | 0.524 | 2.9E-12 | 0.515 | 1.4E-1 |
| *P. tetraurelia* | *P. sonneborni* | 0.137 | 0.269 | 5.5E-12 | 0.515 | 3.7E-1 |
| *P. tetraurelia* | *P. tredecaurelia* | 0.291 | 0.564 | 3.0E-27 | 0.663 | 1.2E-48 |
| *P. tredecaurelia* | *P. biaurelia* | 0.069 | 0.132 | 2.0E-4 | 0.643 | 1.4E-7 |
| *P. tredecaurelia* | *P. decaurelia* | 0.063 | 0.129 | 4.6E-5 | 0.661 | 1.6E-8 |
| *P. tredecaurelia* | *P. dodecaurelia* | 0.059 | 0.117 | 2.7E-4 | 0.664 | 2.3E-8 |
| *P. tredecaurelia* | *P. jennignsi* | 0.142 | 0.197 | 4.5E-2 | 0.528 | 2.1E-1 |
| *P. tredecaurelia* | *P. novaurelia* | 0.044 | 0.094 | 5.4E-4 | 0.655 | 1.9E-5 |
| *P. tredecaurelia* | *P. octaurelia* | 0.070 | 0.170 | 3.6E-9 | 0.614 | 1.7E-5 |
| *P. tredecaurelia* | *P. pentaurelia* | 0.039 | 0.103 | 1.2E-6 | 0.571 | 5.5E-2 |
| *P. tredecaurelia* | *P. primaurelia* | 0.092 | 0.218 | 1.7E-10 | 0.583 | 5.3E-4 |
| *P. tredecaurelia* | *P. quadecaurelia* | 0.094 | 0.233 | 1.2E-11 | 0.704 | 7.2E-17 |
| *P. tredecaurelia* | *P. sexaurelia* | 0.285 | 0.391 | 3.6E-4 | 0.514 | 3.0E-1 |
| *P. tredecaurelia* | *P. sonneborni* | 0.093 | 0.138 | 2.1E-2 | 0.513 | 6.1E-1 |
| *P. tredecaurelia* | *P. tetraurelia* | 0.067 | 0.154 | 1.7E-7 | 0.662 | 2.4E-9 |

**Table S4.** Segmental duplications in Paramecium species.

| species name | Number of segmental duplications | Fraction of retained ohnologs involved in a segmental duplication | Fraction of single ohnologs involved in a segmental duplication | P value |
| --- | --- | --- | --- | --- |
| *P. biaurelia* | 34 | 7.4E-4 | 1.7E-3 | 0.016946 |
| *P. decaurelia* | 26 | 3.5E-4 | 1.7E-3 | 8.45E-05 |
| *P. dodecaurelia* | 31 | 5.1E-4 | 1.7E-3 | 0.000984 |
| *P. jenningsi* | 20 | 5.2E-4 | 1.2E-3 | 0.100879 |
| *P. novaurelia* | 18 | 4.0E-4 | 1.2E-3 | 0.027285 |
| *P. octaurelia* | 21 | 4.2E-4 | 9.7E-4 | 0.086093 |
| *P. pentaurelia* | 278 | 6.5E-3 | 1.1E-2 | 1.30E-05 |
| *P. primaurelia* | 35 | 8.2E-4 | 1.9E-3 | 0.016052 |
| *P. quadecaurelia* | 21 | 4.5E-4 | 1.3E-3 | 0.025925 |
| *P. sexaurelia* | 25 | 4.5E-4 | 1.2E-3 | 0.023424 |
| *P. tetraurelia* | 52 | 1.1E-3 | 2.0E-3 | 0.054674 |
| *P. tredecaurelia* | 54 | 1.2E-3 | 2.4E-3 | 0.015686 |

**Figure S1.**
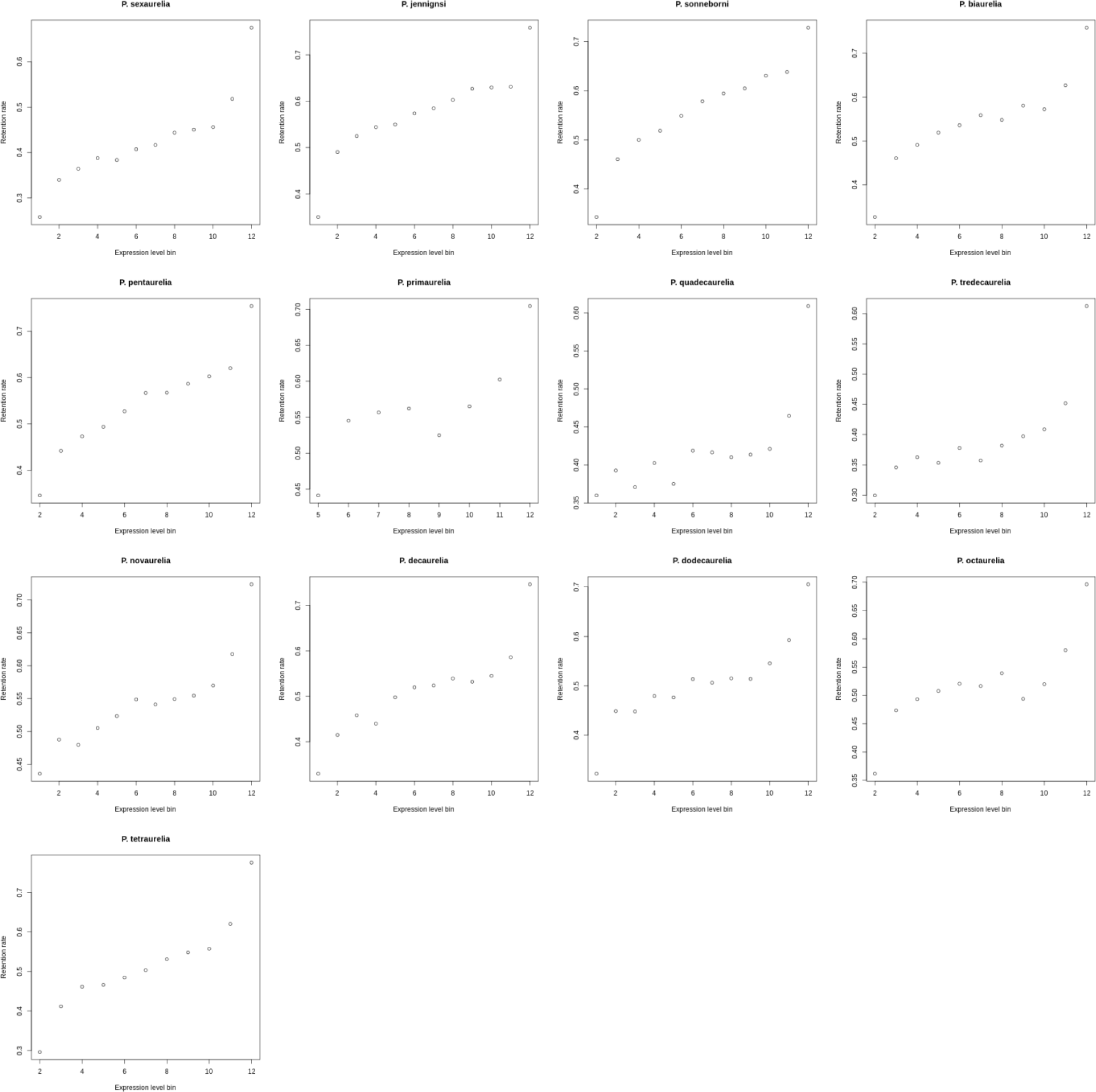
Post-WGD retention rate for genes grouped in bins of expression level (12 bins of equal size) in each *P. aurelia* species.

